# Scalable longitudinal imaging and transcriptomics of cells in dynamic enclosures

**DOI:** 10.64898/2026.05.05.723030

**Authors:** Tarun K Khurana, Liz Y Wu, Pier Federico Gherardini, Seyedsina Moeinzadeh, Mehdi Mohseni, Filiz Gorpe Yasar, Roger Dettloff, Justin Poelma, Shan Sabri, Annarita Scaramozza, Yunmin Li, Nirvan Rouzbeh, Nicholas Elder, Shreya Deshmukh, Alireza Majd, Ria Gupta, Sina Farahvashi, Isaac Thomas, Craig Betts, Frank Charbonier, David Roberts, Pei-Lin Hsiung, Eloise Zargari-Pariset, Richard Yau, Olaia F Vila, Molly Phillips, Carmen Xiao, Jie Wang, Yiqi Zhou, Punam Adhikari, Meng Taing, Elaheh Farjami, Behnam Javanmardi, Merek Siu, The Cellanome Development Team, Camilla Valente, Chris Cox, Kathryn Geiger-Schuller, Shannon J Turley, Orit Rozenblatt-Rosen, Faranak Fattahi, Joseph R Ecker, Jeffrey R Jones, Fred H Gage, Matthew H Spitzer, Jasmine Pritchard, Ilya Kupershmidt, Peter Lundberg, Shawn Levy, Omead Ostadan, Gary P Schroth, Mostafa Ronaghi

**Author notes:** Joint first authors (Tarun K Khurana, Liz Y Wu, Pier Federico Gherardini, Seyedsina Moeinzadeh). ***The Cellanome Development Team***: Reto Schoch, Victor Quijano, Isabel Thomas, Makenzie Sacca, Jessica Finau, Mika Maenaga, Liz Murray, Kian Akhikari, Josh Lamstein, Tim Williams, Maria Johnson, Karl Voss, Paul Kitabjian, Chee Meng Chen, Yiching Siwinski, Matt Hart, Aoran Xu, Abhishek Kurup, Alex Blackburn, Michael Young, Riley McAndrews, Nate Crisostomo, Lena Beir, Julio Costas, Tyler Dill, Mark Fertal, Mailie Bowers, Angel Angeles, Wei Wang, Roya Mazrouei, Sean Wheeler, Jing Men, Chris Chiam, Kit Yeng Wong, Liying Hong, Zhiyi Sun, Harry Vlahos, Shahram Hajizadeh, Yury Frolov, Swanand Valame, John Nguyen, Sarah Connor, MariJo Gallina, Teresa Ai, Kat Chang, Margaret Donovan, Dawn Barry, Praveer Sharma, Aparna Natarajan, Paloma Garcia, Matt Wang, Jerry Naldo, Brian Nunez, Magnolia Bostick, Ouriel Caen, Michael J Schachter, Ali Agah, Niranjan Srinivas.

## Abstract

Dynamic transitions between cell states underlie both normal physiology and disease. However, most single-cell technologies capture only static snapshots. To address this gap, we developed a platform that integrates light-guided hydrogel polymerization with computer vision to generate on-demand compartments around live cells, enabling longitudinal imaging of cellular behavior paired with whole-transcriptome profiling of the same cells at scale.

These data link dynamic phenotypes with molecular programs, enabling deeper characterization of cellular states. This approach revealed an adaptive, drug-resistant state in lung cancer cells characterized by potassium channel upregulation and p53-dependent quiescence. In models of adipogenesis and microglial phagocytosis, joint analysis of imaging and transcriptomic data identified key drivers of cellular function that were missed by transcriptomic clustering alone. These results establish the value of paired functional and transcriptomic analysis to resolve molecular drivers of complex cellular behaviors.

## Introduction

Cell state transitions drive physiological processes as diverse as embryonic development, tissue homeostasis, and immune function, while their dysregulation underlies diseases including cancer, where phenomena like epithelialto-mesenchymal transition promote metastasis(*1*). These transitions involve coordinated rewiring of regulatory networks across epigenetic, transcriptional, and metabolic layers(*2, 3*).

While cell clustering using transcriptomic data has become the gold standard for defining cellular states(*4*), this approach provides only static snapshots that obscure the trajectories, intermediates, and timing heterogeneity inherent in state transitions. For instance, inferring dynamics from population snapshots can be misleading, as they cannot distinguish rapid proliferation from cell cycle arrest(*5*). Fundamentally, transcriptomic analysis can only *infer* cellular function rather than *observe* it.

Here we present a platform that addresses this fundamental gap by enabling longitudinal studies of live cells, directly linking dynamic cellular behavior as observed from imaging to the molecular programs active in each individual cell.

The platform combines advanced computer vision for real-time observation with semipermeable, reversible hydrogel compartments that can be formed by photopolymerization around tens of thousands of individual cells, which we term CellCage Enclosures. This capability effectively creates a massively parallel array of individual wells, allowing scientists to probe, and observe individual cells as they move, signal, differentiate, or respond to stimuli over time. Unlike microwell-based technologies with fixed locations, compartments are dynamically created on-demand. Different from previous approaches(*6* –*9*), we utilize hollow porous hydrogel enclosures and biocompatible materials to enable culture and observation of cells over multiple days or weeks.

At the conclusion of observation, whole transcriptome capture from each enclosure creates an unambiguous link between function, as observed by imaging, and molecular phenotype. Critically, the platform accommodates adherent cells in their attached state and enables studying cellular interactions by including multiple cells per enclosure. The direct pairing of imaging and transcriptomic data, and the ability to provide a substrate for cell adhesion differentiate our approach from recent work based on cell compartmentalization using hydrogel microcapsules in solution(*10, 11*).

This technology enables observing biological processes unfold in real-time and then immediately interrogating the specific molecular drivers of that observed behavior in the exact same cell. We demonstrate that the resulting datasets combining simultaneous imaging and transcriptomics of individual cells represent a fundamentally new datatype that enables characterization of cell states to a level of detail not possible with transcriptomics alone.

## Results

### Highly parallel compartmentalization of target cells

Our platform employs light-guided hydrogel polymerization to create compartments on-demand called CellCage Enclosures (CCEs) around target cells (Fig. 1). The workflow is automated on an instrument combining advanced multi-wavelength fluorescence imaging, Digital Micromirror Device (DMD) projection, GPU-based computation, and automated fluidics.

**Figure 1:**
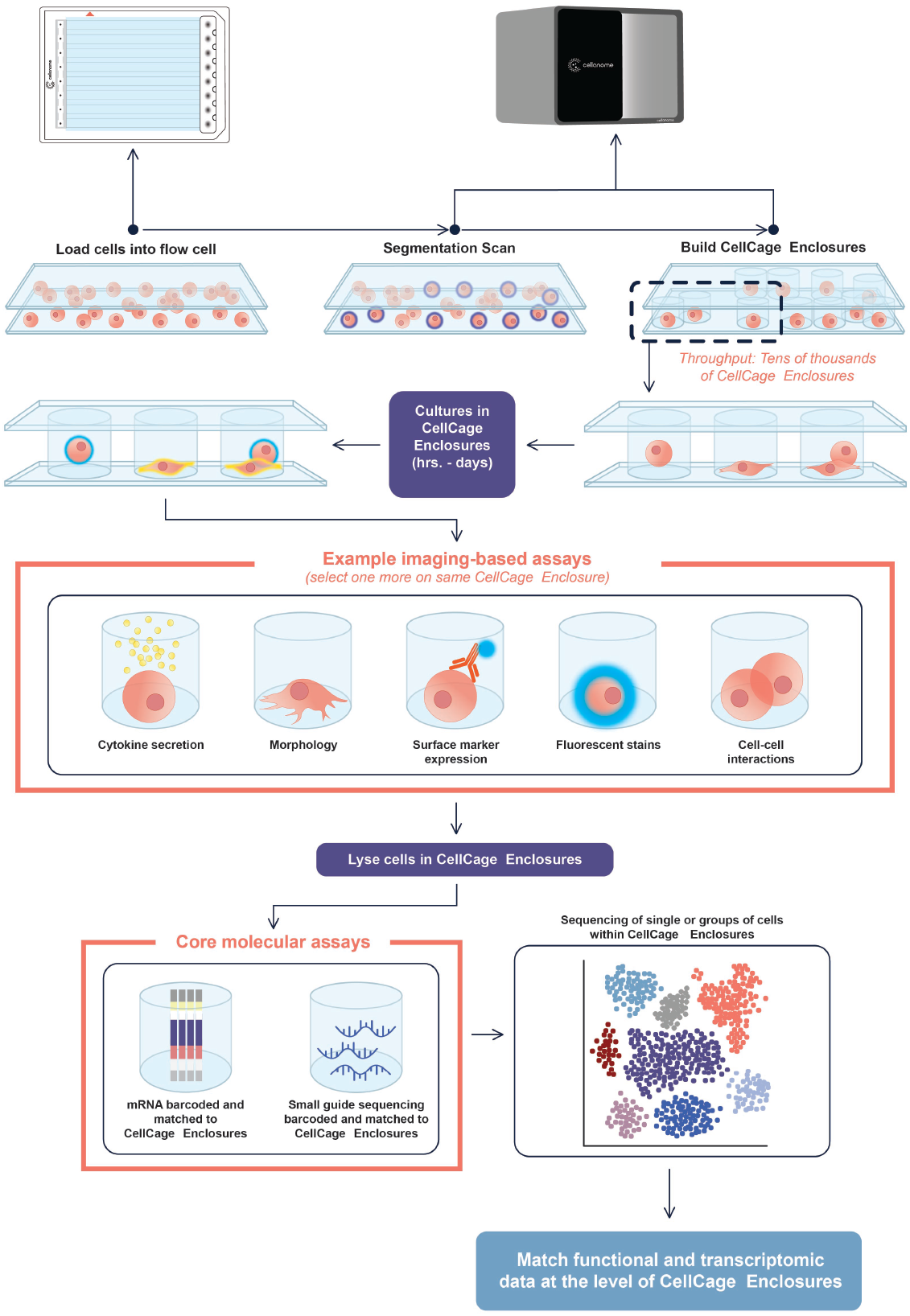
Overview of the workflow. The flow cell has the same footprint as a microtiter plate and features eight individually addressable lanes, ∼75 µm in height. Cells are mixed with a hydrogel precursor, loaded in a lane, and imaged. The position of the cells is then identified using computer vision, and the optimal placement of CellCage Enclosures (CCEs) is calculated. A Digital Micromirror Device (DMD) is then used to project light in a pattern corresponding to the desired location and shape of the CCEs within the field of view. The light activates a photo initiator, which induces polymerization of the hydrogel thus forming the CCEs within a few seconds. This procedure is then repeated for all the fields of view that comprise the flow cell lane. Reagents can be delivered, and media can be exchanged, at user defined time points, and compartmentalized cells can be serially imaged over several days to perform a variety of assays. Finally, cells can be lysed to prepare sequencing libraries that can be matched to individual CCEs. Representative assays are listed; the platform is flexible and can support additional imaging- and sequencing-based assays.

Cellular suspensions are first mixed with hydrogel precursor and a photoactivator prior to loading into an eight-lane flow cell (with the same footprint as a microtiter plate). Each 9 cm × 6 mm lane supports independent assay parameters, including different media, drug treatments, etc. Computer vision is used to identify the position of each cell, followed by calculation of a digital photomask representing optimal CCE placement around individual cells or cell groups (depending on settings) to maximize CCE density while preventing overlap. The DMD is used to project the photomask with 405nm light, photopolymerizing hollow hydrogel compartments around target cells with the glass surfaces of the flow cell forming the top and bottom. A single lane is processed in under 15 minutes at 4× magnification. Unpolymerized precursor is washed away and replaced with cell culture media, leaving compartmentalized cells ready for live-cell analysis.

CCEs offer flexible geometry and size configurations. A single lane can accommodate 5,000-8,000 CCEs with 90 µm diameter, with the exact throughput depending on the size of the CCE, the sample density and the uniformity of cell loading (Supplementary Fig. 1). Larger CCEs (400 µm diameter) enable compartmentalization of hundreds of complex cell aggregates per lane (e.g. neurospheres, Supplementary Fig. 2). Moreover, the compartmentalization logic enables the formation of CCEs containing multiple cells, which can be used to study biological processes that depend on cellular interactions (Supplementary Fig. 3). The permeability of the walls can be tuned to allow exchange of small molecules and proteins for dynamic reagent delivery, diverse culturing protocols and staining with antibodies (Supplementary Fig. 4).

The biocompatible hydrogel system provides fast gelation (seconds), degradability, and tunable porosity supporting diffusion of molecules in a 5-2000 kDa range, enabling cell culture over weeks with optional media exchange (Fig. 2A-D). Orthogonal gel chemistries, which can be degraded by reducing versus oxidizing agents, specific enzymes or light, enable sophisticated workflows including enrichment and elution of specific sub-populations of interest based on multi-functional characteristics, facilitating the automation of complex biological assays without requiring separate instrumentation (Fig. 2E-F).

**Figure 2:**
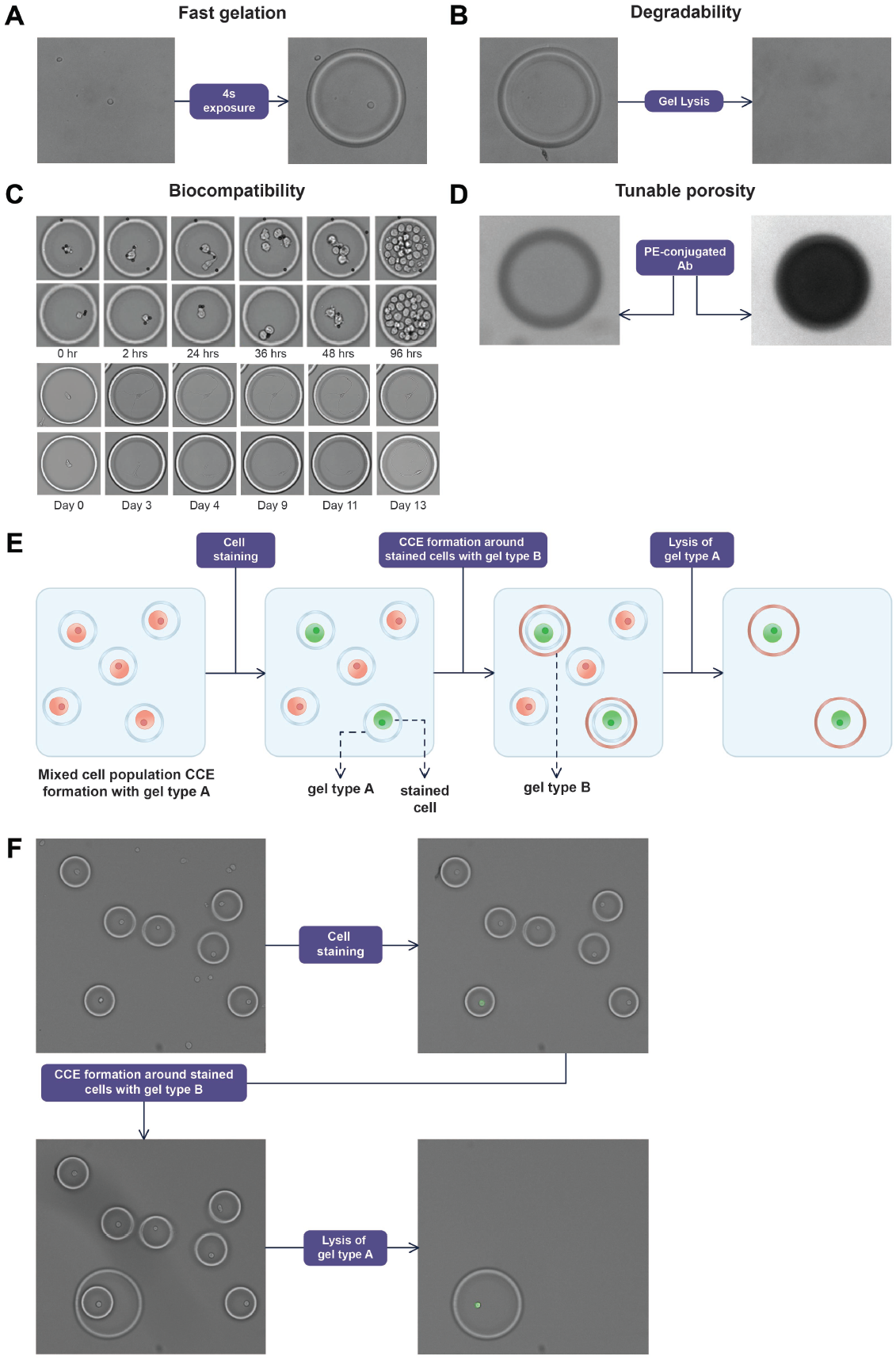
Key properties of the hydrogel material that enable manipulation and longitudinal observation of cells. A: fast gelation following light exposure. B: degradability. C: biocompatibility. The top two rows depict primary CD8 T cells cultured over a period of 4 days in the presence of CD3/28 activation beads. The bottom two rows depict induced neurons (61) cultured over 13 days on a flow cell coated with PLO/Laminin. D: tunable porosity. The picture depicts different degrees of CCE permeability to a PE-conjugated antibody. The CCE on the left is permeable to the fluorescent antibody, while the one on the right is not. E: diagram of the selective retention workflow using hydrogels with orthogonal degradation properties. F: selective retention of NK92 cells from a mixture of Jurkat and NK92 cells. NK92 cells were labeled using a PE-Conjugated anti-CD56 antibody.

### Transcriptome measurements in CellCage Enclosures

To integrate longitudinal cellular phenotyping with genome-wide transcriptomics, we developed flow cell surfaces spotted with mRNA capture oligonucleotides containing approximately 23,000 unique capture spots per lane to unambiguously assign transcriptomes to individual CCEs (Fig. 3A). Each capture spot contains identical oligonucleotides with a universal priming site and a polyT handle for 3^*′*^ mRNA capture and cDNA synthesis.

**Figure 3:**
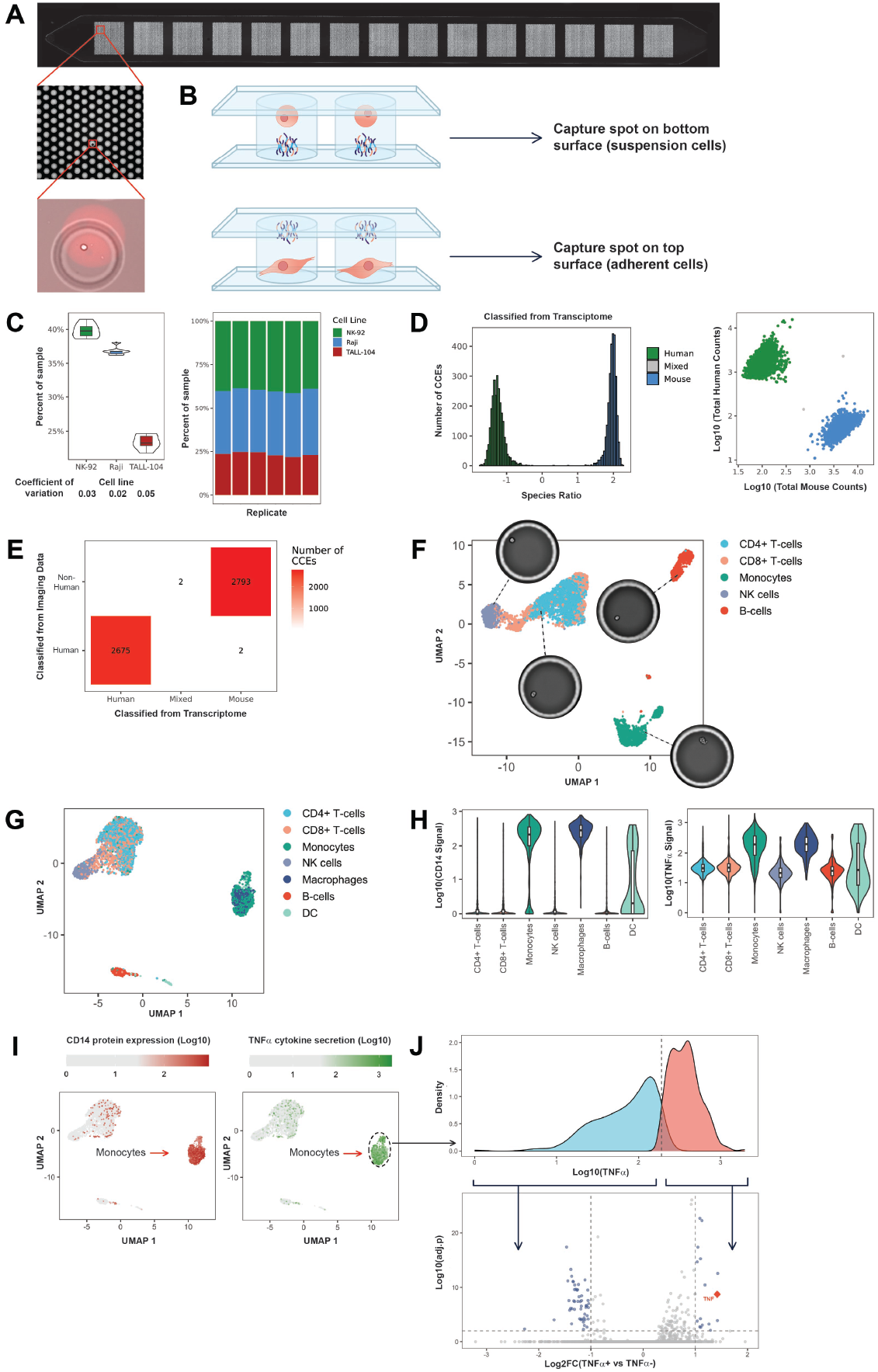
Transcriptome measurements in CellCage Enclosures. A: surface with capture spots for transcriptomic assays. The insets display different levels of magnification, with the top image representing an entire flow cell lane which contains ∼23,000 unique capture spots arranged in 15 separate fields, the middle representing a portion of a field with 140 individual capture spots visible, and the bottom displaying a single CCE comprising a cell and a portion of an individual spot. B: for analysis of suspension cells, we use a flow cell where the bottom surface is spotted with capture oligos. For adherent cells the bottom surface is coated with proteins such as fibronectin and laminin for cell adhesion, and the capture oligos are spotted on the top surface. C: To assess the technical reproducibility of the transcriptome assay, we used a sample containing a mixture of three different cell lines (NK-92, Raji, TALL-104) that was compartmentalized and processed in parallel across six different flow cell lanes. The violin plots on the left represent the estimated proportions (y axis) of different cell lines (x axis) from transcriptomic signatures (see Methods) in a mixed sample across six technical replicates on six different lanes of a flow cell. The numbers under the x axis represent the coefficients of variation. The stacked bar plots on the right represent the estimated proportion of the different cell types in each lane. D: mixed species (Barnyard) experiment to assess the specificity of transcriptome capture. A mixture of EG7-OVA (mouse) and Jurkat (human) cells were compartmentalized as individual cells. Each CCE was assigned to a species based on the log-ratio of the reads from each species. The histogram on the left shows the distribution of species ratio values across the CCEs, colored by the species assignment. The scatterplot on the right shows the counts of human (y axis) and mouse (x axis) transcripts. Each dot represents an individual CCE, colored by species assignment. E: in the Barnyard experiment the Jurkat cells constitutively express an RFP reporter, enabling their identification by imaging. The plot shows the confusion matrix between the imaging-based (y axis) and transcriptomic-based (x axis, see panel D) classification of each CCE. F: UMAP displaying the results of a transcriptomic assay on healthy human PBMC. The colors correspond to cell type labeling based on the expression data, using the SingleR(62) package with the signatures from the Blueprint ENCODE project(63), which resulted in the expected proportions of major cell types. The pictures represent single cells in CCEs that were mapped to each cell type. G: UMAP based on transcriptomic data from an experiment comprising concurrent measurement of CD14 surface protein expression and TNFα cytokine secretion paired with transcriptome in healthy human PBMC stimulated with LPS. The points are colored by predicted cell type using SingleR with the signatures from the Blueprint ENCODE project. H: violin plots of CD14 expression (left) and TNFα secretion (right) in different cell types annotated from the transcriptomic data (x axis). I: Overlay of CD14 surface protein expression (left) and TNFα cytokine secretion (right) on the UMAP derived from the transcriptomic data. J: the distribution of TNFα cytokine secretion in monocytes was used to define TNFα-positive and -negative populations. The volcano plot shows the results of the differential gene expression analysis between the two populations.

The CCE formation algorithm ensures each compartment overlaps one or more capture spots, while restricting each capture spot to a single CCE for unambiguous read assignment. For suspension cells, capture spots are printed on the bottom surface; for adherent cells requiring surface coatings, capture spots are spotted on the top surface, allowing the bottom to be treated with surface coating materials to promote cell adhesion (Fig. 3B). Importantly, coated flow-cells enable analysis of adherent cells in a physiologically relevant state, which is not possible with droplet-based technologies.

We validated the reproducibility and specificity of the assay (Fig. 3C-E) and demonstrated its ability to resolve immune cell populations present in peripheral blood mononuclear cells (PBMCs), where we obtained 3782 median transcripts and 1338 median genes at a depth of ∼27,000 reads per CCE (Fig. 3F). By combining transcriptional analysis with surface protein staining and cytokine detection measurements we observed specific secretion of TNF*α* by CD14^+^ monocytes following stimulation of healthy human PBMCs with LPS (Fig. 3G-J), matching previous observations(*12*).

### Integrated sequencing and imaging, a new datatype for deeper cell characterization

The integration of live cell imaging of adherent and suspension cells with transcriptomic readouts at scale represents a fundamentally new datatype. While imaging data can be quantified by conventional instance segmentation and feature extraction, we sought to leverage the DINOv2 image foundation model(*13*) to extract rich morphological descriptors from cellular images, with dimensionality comparable to gene expression readouts.

We validated that 1024-dimensional DINOv2 embeddings can be used to cluster cell lines according to their morphology (Fig. 4A) and are robust to rotation (Fig. 4B). We then integrated these image-derived features with transcriptomic data to characterize the process of adhesion of Hs675.T colon fibroblasts on a fibronectin-coated flow cell (Fig. 4C).

**Figure 4:**
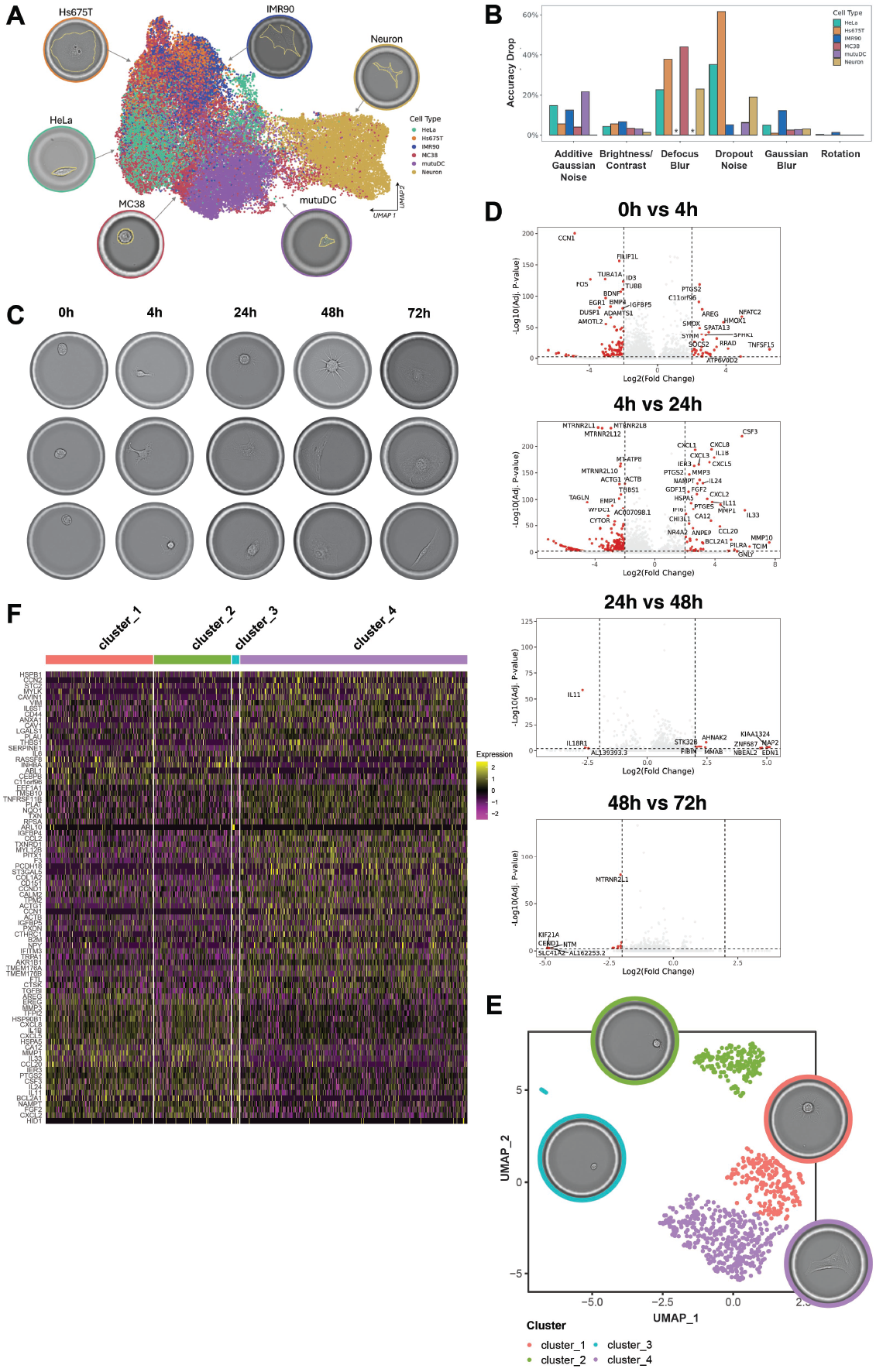
Integrated sequencing and imaging, a new datatype for deeper cell characterization. A: UMAP representing 1024-dimensional DINOv2 features from six cell lines showing clustering by cell identity, confirming capture of meaningful morphological differences. B: effect of common image impairments on the prediction accuracy of a linear classifier trained to predict cell identity based on DINOv2 features (see Methods). Different types of impairments were used (x axis) and the bar plot shows the drop in classifier performance (y axis) for each impairment and cell line (identified by the color). The stars denote two cell lines for which defocused blurred images were not available. Importantly, random rotations result in negligible drops in accuracy, which is key to the use of these morphological features, as cell position cannot be controlled. C: morphological appearance of Hs 675.T colon fibroblast grown on a flow cell with a fibronectin-coated bottom surface, and a top surface with capture spots for transcriptomic analysis. Note that in this experiment we performed transcriptomic analysis at each timepoint in different lanes, which requires cell lysis. Therefore, in this case the pictures depict representative images at each time point, not longitudinal images of the same cells. D: volcano plots displaying the results of pseudo-bulk differential expression analysis between consecutive timepoints. E: UMAP visualization and clustering of 1024-dimensional embeddings extracted by DINOv2 applied to individual cell images at the 24 hours timepoint. The pictures display representative images of each cluster. F: single-cell differential expression analysis between the cell morphology-derived clusters identified in panel E.

The 24 hours timepoint represents a critical transition in the culture when proliferative programs become downregulated and pathways mediating response to cytokines known to mediate fibroblast differentiation (IL-1, IL-4, IL-10, IL-13) became prominent (Fig. 4D and Supplementary Fig. 5).

We examined images for 834 CCEs after 24 hours of culture and observed a diversity of morphological appearances reflecting different stages of adaptation to the culture in the unsynchronized cells. Using DINOv2 to extract embedding vectors capturing visual features, followed by UMAP visualization and K-means clustering, we identified four morphotypes: cluster 3 (spherical, non-adhered), clusters 1-2 (spherical with thin spreading processes), and cluster 4 (fully adhered) (Fig. 4E).

Differential expression analysis between the morphotypes revealed distinct transcriptional programs with fully adhered cells expressing higher levels of adhesion molecules (PCDH18, LGALS1), collagen (COL1A2), connective tissue growth factor(*14*) (CCN2), cytoskeleton proteins (TMSB10), and mature fibroblast markers (F3(*15*), PLAT (*16*)), along with CCL2 that mediates survival signaling(*17, 18*) (Fig. 4F). Partly adhered cells expressed higher level of extracellular matrix remodeling enzymes (MMP3, MMP1) and FGF2. They also expressed a different set of cytokines and chemokines including CCL20, which is downstream of IL-1*β*/TGF-*β* signaling(*19*), and IL-33, which is upregulated in Ulcerative Colitis in both murine models(*20*) and patient samples(*21, 22*).

These results demonstrate that quantitative morphological features from brightfield imaging correspond to discrete cellular states with markedly different gene regulatory programs, even in supposedly homogenous cell cultures, a dynamic that is obscured in bulk measures or droplet-based approaches incompatible with cellular adherence.

### Longitudinal imaging combined with transcriptomic identifies drug resistant states

Longitudinal imaging provides key phenotypic insights that cannot be resolved by individual snapshots, and it is particularly powerful when combined with transcriptomic readouts. We sought to demonstrate this unique feature of our platform by imaging A549 lung cancer cells treated with Olmutinib, an EGFR tyrosine kinase inhibitor, every eight hours over 32 hours, followed by cell lysis and 3^*′*^ mRNA sequencing. The final dataset comprised 10,604 treated and 4,337 untreated CCEs. Differences between treated and untreated cells were as expected (Supplementary Fig. 6), and subsequent analysis focused on the heterogeneity within the treated condition.

To resolve the relationship between transcriptomic states and cellular behavior, we classified individual treated CCEs into four imaging-derived phenotypes (Fig. 5A): *cell cycle arrest, proliferation, dead*, and *daughter cell resistant*—the latter representing a striking phenotype where cells divided under drug pressure with one daughter dying while others persisted. Interestingly, these phenotypes did not strongly segregate with cell clusters defined from the transcriptional data alone (Fig. 5B-C), as all clusters contained representatives from all imaging-derived phenotypes (Fig. 5D).

**Figure 5:**
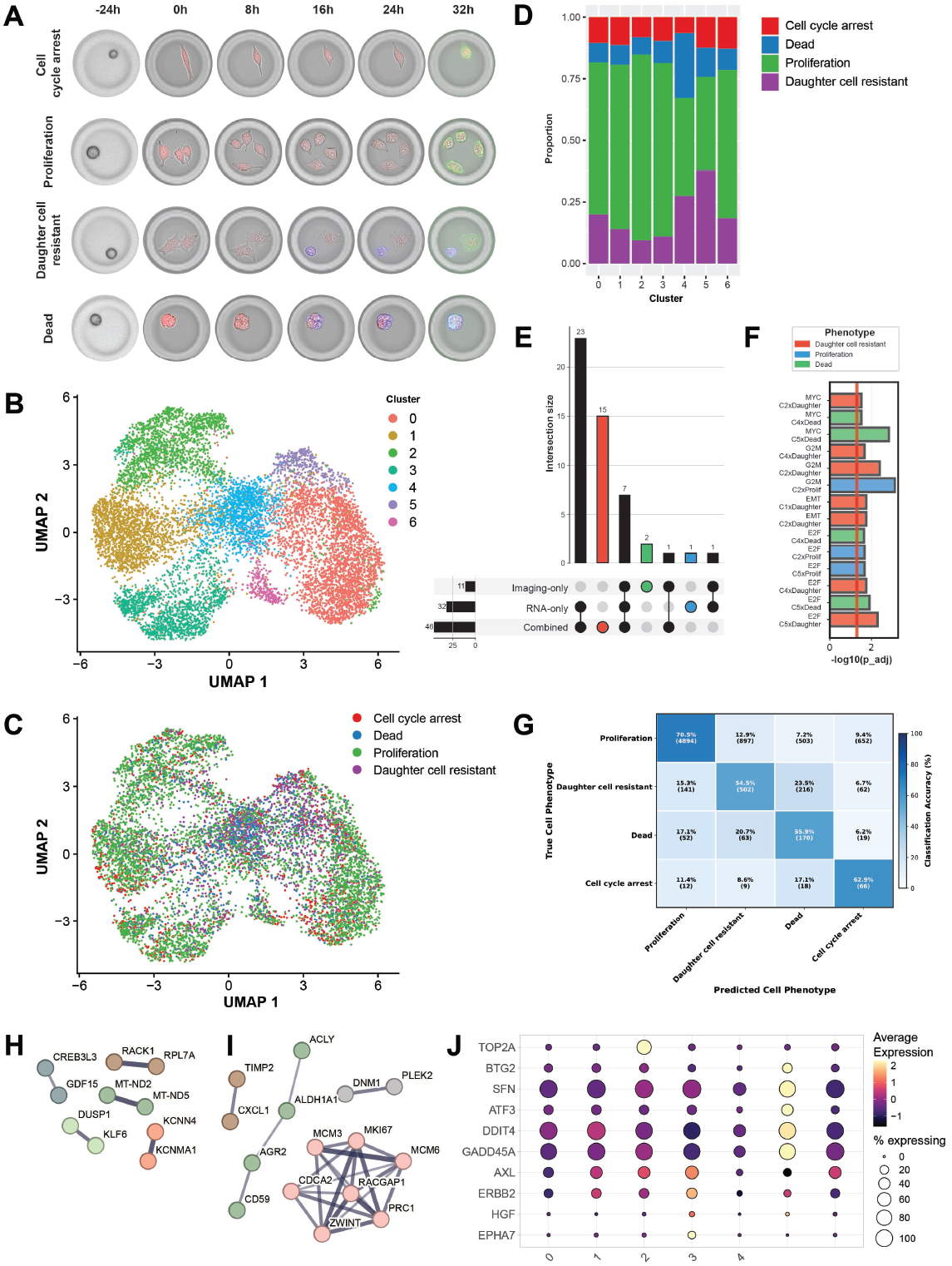
Profiling A549 cells following treatment with Olmutinib. A: classification of CCEs in different phenotypes based on the analysis of longitudinal imaging data. Red: CellTrace™ Far Red, blue: Annexin V, green: EGFR. B: UMAP based on the transcriptomic data from 10,604 CCEs containing A549 cells treated with 10 µM Olmutinib. The colors represent different transcriptomic clusters. C: UMAP based on the transcriptomic data (same as panel B) colored according to the imaging-derived CCE classification in panel A. 2,328 CCEs that could not be accurately classified were excluded from the analysis. D: proportion of CCEs (y axis) belonging to each imaging-based phenotype (indicated by the color) within each gene expression cluster (x axis). E: Upset plot showing overlap of significant GSEA pathway enrichments across three classification strategies. The combination of transcriptomic clustering with imaging classification identified 15 unique pathways not found in either single-modality strategy. F: significant interaction effects (p_adj < 0.05) between RNA clusters and imaging phenotypes on the prediction of drug resistance pathway modules (G2M checkpoint, E2F targets, MYC targets, DNA repair, EMT) (see Methods). The daughter cell resistant phenotype showed 7 out of 14 total significant interactions, indicating that pathway activities are maximally explained by the combination of transcript state and the daughter cell resistant phenotypic classification. G: Confusion matrix for elastic net prediction of imaging phenotypes from gene expression. H: STRING PPI network for top 50 positive coefficient genes (associated with daughter cell resistance). I: STRING PPI network for top 50 negative coefficient genes (inversely associated with daughter cell resistance). J: Selected differentially expressed genes between expression-defined clusters (x axis). The color represents the average expression (scaled per gene) and the size of the circle indicates the percentage of CCEs expressing the gene. Cluster 2 showed strong enrichment for cell division pathways and overexpressed the proliferation marker TOP2A. Cluster 3 exhibited activation of multiple EGFR bypass pathways with overexpression of EPHA7 (64), HGF (65), ERBB2 (66), and AXL(67), all capable of activating MAPK signaling independently of EGFR. Cluster 5 displayed enrichment of p53 targets, including upregulation of quiescence-associated genes such as GADD45A, REDD1, ATF3, SFN, and BTG2.

When individual CCEs were classified by combining the imaging phenotypes and gene expression-based clusters, we identified 46 significant pathway enrichments, including 15 not found when using either modality alone (Fig. 5E). Regression analysis revealed that combining data modalities predicts drug resistance pathway modules (including G2M/E2F/MYC activities) significantly better than either data type alone (Fig. 5F). The daughter cell resistant phenotype showed the highest number of significant interactions, supporting the notion that it represents a distinct regulatory state.

To identify key genes driving phenotypic differences, we developed an elastic net model to classify CCEs into imaging-derived phenotypes using expression data, achieving 68% overall accuracy (Fig 5G). Focusing on molecular drivers of the daughter cell resistant phenotype, the top-50 positive and negative coefficient genes from the elastic net model were used to infer protein-protein interaction networks using STRING(*23*). Positive predictors (genes associated with resistance) included mitochondrial-encoded genes (MT-ND2, MT-ND5), transcription factors (KLF6, CREB3L3), stress response genes (DUSP1, GDF15), and calcium-activated potassium channels KCNMA1 and KCNN4 (Fig. 5H). KCNN4 expression is associated with chemoresistance in multiple cancer types(*24, 25*) and its blockade in erlotinib-resistant A549 cells can overcome resistance(*26*). Moreover, both KCNMA1 and KCNN4 are activated by oxidative stress responses in melanoma(*27*).

Negative predictors formed a tightly connected network of cell cycle machinery genes: MCM6, MKI67, ZWINT, PRC1, RACGAP1, CDCA2, and MCM3 (Fig. 5J). The daughter cell resistant phenotype was enriched in transcriptomic cluster 5, which displayed higher expression of p53 targets (Supplementary Fig. 7), including upregulation of quiescence-associated genes such as GADD45A(*28*) (G2/M arrest), REDD1(*29*) (stress-induced modulator of mTOR), ATF3(*30*) (quiescence and chemoresistance), SFN(*31*) (G2/M inhibition), and BTG2(*32*) (quiescence regulation) (Fig. 5K). Since A549 cells are p53 wild-type, these findings support the notion that intact p53 function promotes persistent, slow-cycling, drug-resistant states that confer resistance to targeted therapies and enhanced invasive capacity(*33* –*37*).

These results point to the daughter cell resistant phenotype representing a specific cellular state associated with resistance characterized by upregulation of potassium channels and a quiescent/slow-cycling phenotype enabled by p53. This state, which is uniquely identified by the ability to observe individual cells over time, reveals that sibling cells originating from the same progenitor can adopt divergent fates under drug pressure, in a process that does not depend on the acquisition of new mutations. This phenomenon does not represent selection of a pre-existing state and is in accordance with theoretical models of cellular adaptation based on stochastic sampling of gene expression space(*38* –*40*).

### Multi-modal data enables the development of supervised models that identify key transcriptional drivers of cellular function

Finally, we investigated the extent to which transcriptomics can predict complex cellular functionality by studying lipid droplet accumulation in differentiating preadipocytes and microglial phagocytosis, using fluorescence imaging as ground-truth to define cellular functions.

We cultured primary human preadipocytes for seven days on a fibronectin-coated flow cell, in the presence of differentiation media. Cells were stained with a nuclear stain (Hoechst) and BODIPY to measure lipid content followed by lysis and 3^*′*^ mRNA sequencing. The dataset comprised 673 CCEs, each containing a single cell at the beginning of the experiment. Transcriptomic clustering failed to identify adipocytes with high-lipid content, revealing that genes driving global gene expression differences are not necessarily those controlling cellular functions of interest (Fig. 6A-C). Using elastic net modeling with lipid-to-nuclear stain ratios as the response variable (Fig. 6D), we identified 85 predictor genes, only two of which overlapped with top-20 cluster-specific differentially expressed genes (Fig. 6E). Moreover, differential expression fold-changes between transcriptomic clusters showed no correlation with model coefficients (Fig. 6F), and the top-3 positive predictors showed broad distribution across the UMAP space defined by the totality of the gene expression data (Fig. 6G). These results reinforce the value of paired functional and transcriptional data to elucidate key molecular drivers of cellular function.

**Figure 6:**
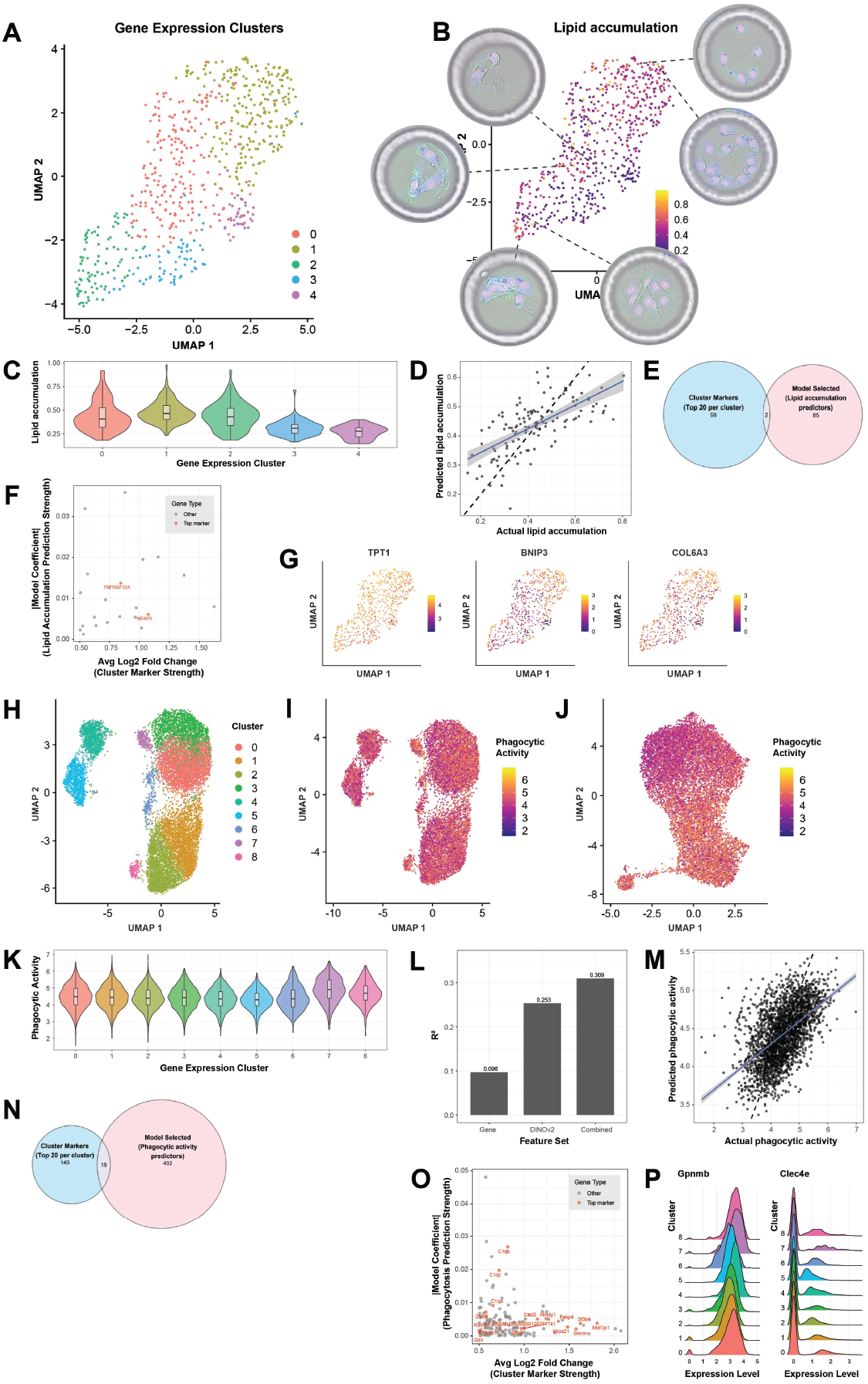
Unsupervised clustering of transcriptomic data fails to capture complex cellular phenotypes, including lipid accumulation by preadipocytes and phagocytic activity of microglial cells. A: UMAP based on transcriptomic data from primary human preadipocytes differentiated for seven days on a fibronectin-coated flow cell. The colors correspond to different clusters based on transcriptomic analysis. B: Transcriptomic UMAP colored by the lipid accumulation score, defined as the ratio between the BODIPY stain and the nuclear stain in each CCE. The insets show examples of cells that are very close in gene expression space but differ in their lipid content. C: violin plots depicting the distribution of lipid accumulation scores (y axis) across the transcriptomic clusters (x axis). D: actual (x axis) vs predicted (y axis) lipid accumulation scores from the elastic net model. The plot is for the held-out test set (20% of the total data). E: Euler diagram showing the overlap between top-20 differentially expressed genes between transcriptomic clusters (blue) and model-selected predictors of lipid accumulation (pink). F: average Log2 fold-change between clusters (x axis) vs absolute model coefficient (y axis) for the genes selected by the model. The red color indicates genes that are among the top-20 differentially expressed genes between transcriptomic clusters. G: Gene expression UMAP colored by the top-3 positive predictors identified by the model, showing that the expression values of these genes are uniformly distributed across the UMAP based on global transcriptomic differences. H: UMAP based on transcriptomic data for BV2 mouse microglial cells. The colors correspond to different clusters based on transcriptomic analysis. I: transcriptomic UMAP colored by phagocytic activity as measured by pHrodo™ intensity after four hours. J: UMAP based on DINOv2 features, colored by phagocytic activity showing a greater degree of separation between high vs low phagocytic scores, compared to the transcriptomic UMAP in panel H. K: violin plots depicting the distribution of phagocytic scores (y axis) across the transcriptomic clusters (x axis). L: R^2^ performance of elastic net models trained on expression-only features, DINOv2-only features or a combination of the two (x axis). The data refers to the held-out test set (20% of the total data). M: actual (x axis) vs predicted (y axis) phagocytic scores from the elastic net model using the combined expression and DINOv2 features. The plot is for the held-out test set (20% of the total data). N: Euler plot showing the overlap between top-20 differentially expressed genes between transcriptomic clusters (blue) and model-selected predictors of phagocytic activity (pink). O: average Log2 fold-change between clusters (x axis) vs absolute models coefficient (y axis) for the genes selected by the expression-only model. The red color indicates genes that are among the top-20 differentially expressed genes between transcriptomic clusters. P: ridge plots displaying the expression level (x axis) of Gpnmb and Clec4e across transcriptomic clusters (x axis). These two genes are among the top positive predictors for the gene expression-based model and have clear mechanistic evidence linking them to the phagocytosis process. However, their expression is very similar across all the transcriptomic clusters.

The model identified coordinated pathways necessary for adipocyte differentiation including extracellular matrix remodeling(*41*) (COL6A3, COL6A2, COL3A1), regulators of lipid droplet biogenesis (CIDEC/FSP27, LPIN1), cytoskeleton-associated proteins (TPT1(*42*) – top positive predictor, TMSB10, INF2), mitochondrial proteins (BNIP3, TOMM70, MRPL35, ISCA2, TRAP1) and DNA repair pathways (XPA, RECQL, NSMCE1) as predictive of high lipid content. While cytoskeleton remodeling during lipid accumulation is documented(*43, 44*), the involvement of TMSB10 and INF2 in adipogenesis has not been reported. Similarly, while isolated reports document the association of DNA repair pathways with adipogenesis(*45* –*47*), the genes identified here have not been previously implicated. Negative predictors included genes with reported inhibitory effects on preadipocytes differentiation and lipid accumulation (e.g. TNFRSF12A (*48*), TRIO(*49*), SKP1(*50*), CCN2(*51*), SERPINA3(*52*)), as well as multiple novel candidates (MRFAP1L1, MPHOSH8, ARL5A).

To study the extent to which transcriptomics can predict phagocytic activity, we loaded mouse BV2 microglial cells on a fibronectin-coated flow cell in the presence of pHrodo particles, which fluoresce within the acidic environments of endosomes and lysosomes. After four hours the cells were imaged and lysed for transcriptomic anal-ysis, yielding 12,972 CCEs. Cell clusters based on gene expression data did not correlate with phagocytic activity, as measured by pHrodo staining (Fig. 6H-I,K), while DI-NOv2 brightfield image embeddings had better correlation (Fig. 6J). We built elastic net models using gene expression, image embeddings, or both to predict phagocytic activity. Morphology alone outperformed gene expression. The combined model performed best and included 617 features in total, with 15% coming from morphology data and 85% from gene expression data, demonstrating the richness of paired analysis (Fig. 6L-M). Again, we observed poor overlap between model predictors and cluster-specific differentially expressed genes (Fig. 6N), and no correlation between fold-changes and model coefficients for the model using only gene expression data (Fig. 6O).

The top positive predictors in the gene expression model included Gpnmb(*53, 54*) and Clec4e(*55*), two genes for which there is clear mechanistic evidence for their direct involvement in this process, though neither gene was in the top-20 cluster-specific genes, and their expression was very similar across all transcriptional clusters (Fig. 6P). The top negative predictors included the complement genes C1qb and C1qc, striking given their known role in microglial phagocytosis(*56, 57*), suggesting that the cells with highest phagocytic activity did not express the highest levels of complement, mirroring similar observations relating cytokine secretion and killing activity in individual T(*58*) and NK(*59*) cells.

These findings demonstrate that pairing imaging and gene expression data in individual cells enables discovery of genes driving cellular functions of interest by leveraging the intrinsic variability of the biological system, highlighting the value of concurrent multi-modal data. Crucially, these discoveries were not possible by transcriptomic analysis alone.

## Discussion

The platform presented here overcomes the challenge of linking dynamic cellular functions to end-point transcriptomics in the same individual cells at scale. By using photoactivatable hydrogels to create on-demand CellCage Enclosures, our system provides the first platform, to our knowledge, capable of acquiring longitudinal, multi-modal data for tens of thousands of live cells in a single experiment, each paired with end-point gene expression measurements.

Paired longitudinal live cell imaging and transcriptomics in the same individual cells constitutes a fundamentally new datatype, with a unique ability to characterize cellular states and identify molecular drivers of cellular function, as demonstrated in multiple applications. Within the context of the development of virtual cell models, it has been suggested that single timepoint scalar measurements (like fluorescence intensity or cell counts) do not capture enough information to fully map phenotypic space. Instead, high-content, time-resolved imaging data is key to understanding how genetic perturbations influence cellular behavior, especially given the dynamic and multifaceted nature of cellular responses(*60*).

A central advantage is the compatibility of the platform with adherent, large, and fragile cells like neurons, which are often not well-suited for droplet encapsulation or flow cytometry. The customizable flow-cell surfaces allow adherent cells to be analyzed in a state that better preserves their native physiology, eliminating the impact of enzymatic detachment or other manipulations that can alter gene expression. Furthermore, the ability to construct enclosures containing multiple cell types unlocks new opportunities to study phenotypes that depend on cellular interactions (e.g., in neurology, oncology, and immunology), and their heterogeneity as a function of cell ratios.

The system is highly flexible, as nearly every aspect of CCEs can be customized, including size, porosity, shape, and top and bottom surface functionalization, thus enabling the development of a wide variety of live-cell functional assays. More broadly, the core technology, which combines computer vision, photochemistry, and high-density hardware, constitutes a general-purpose automation platform. It enables the development of complex experimental workflows with real-time control feedback at a density far exceeding that of conventional laboratory robotics. Work is underway to expand these capabilities and further automate workflows on the platform.

In conclusion, we developed a technology that directly links the life history and functional behavior of a cell to its molecular state. By enabling the resolution of complex relationships between genome, transcriptome, and cellular behavior, this platform provides a transformative tool for biology and empowers a deeper, more causal understanding of cell states and functions.

## Materials and Methods

### Tissue culture

Human healthy PBMC were purchased from AllCells. PBMC were rested in RPMI-1640 medium supplemented with GlutaMax (Fisher scientific, #31-980-030), 10% Fetal Boving Serum (FBS) (ATCC) and 1% Pen-Strep (Sigma).

Jurkat cells were purchased from ATCC (TIB-152) and cultured in RPMI 1640 medium (Gibco) supplemented with 10% FBS (Gibco) and 1% Antibiotic-Antimycotic 100x (Gibco).

Human Natural Killer (NK-92) cells were purchased from Creative Biolabs and cultured in 75% SuperCult Alpha MEM (Creative Bioarray), supplemented with 12.5% heat inactivated FBS (Creative Bioarray), 12.5% PremaSera Horse Serum (Creative Bioarray), 2 mM L-glutamine, 36 ug/mL Myo-inositol (Sigma 17508), 1x PenStrep (Sigma), 55 uM 2-mercaptoethano (Gibco), 22 uM Folic acid, and, and 200 U/mL IL-2.

MutuDC cells were purchased from Abm (T0528) and cultured in IMDM medium (Gibco), supplemented with GlutaMAX, 10% heat-inactivated FBS, 50uM Beta mercaptoethanol, 1% Pen-Strep.

Hs675.T cells were purchased from ATCC (CRL-7400) and cultured in DMEM (Fisher scientific) supplemented with Insulin (5 µg/mL), FGF-2 (10 ng/mL) and 1% Pen-Strep.

Raji cells were purchased from ATCC (CCL-86) and cultured in RPMI 1640 medium supplemented with 20% FBS and 1% Pen-Strep. TALL-104 were purchased from ATCC (CRL-11386) and cultured in RPMI 1640 medium supplemented with 20% FBS, 1% Pen-Strep and 200 U/mL IL-2.

Human lung cancer A549 cells were purchased from ATCC (CCL-185), and cultured in DMEM supplemented with 10% FBS and 1% Pen-Strep.

Mouse BV2 microglia cells were purchased from Accegen (#ABC-TC212S) and cultured in DMEM supplemented with 10%FBS and Glutamax.

Induced neurons were prepared as described previously(*61*).

### Image analysis

Two different Deep Learning models are used for image analysis. For CCE formation we use the Ultralytics (www.ultralytics.com) implementation of YOLOv5(*68*), a fast detector model that provides bounding boxes of the objects of interest. For data analysis after the completion of the experiment we use the pytorch (pytorch.org) implementation of Mask R-CNN(*69*), which provides detailed masks of each object instance. Both models were trained with an in-house dataset consisting of ∼3 million manually labelled object masks across ∼15,000 images. Segmentation is usually done on brightfield images but can also be done on single-channel fluorescent images for CCE formation. Quantification of imaging data for analysis is typically done by calculating the mean pixel intensity per imaging channel, across all the pixels that comprise an object instance. Background subtraction is performed by subtracting from this value the mean intensity of all the pixels within a CCE that do not belong to any object. To extract unsupervised morphology features we used the DINOv2 large model(*13*) on individual cell crop images. The image analysis pipeline was written using python and Nextflow(*70*).

### CCE formation and characterization

The macromonomer solution is mixed with a photo-initiator solution at a 3:2 volume ratio to make a hydrogel precursor solution. The hydrogel precursor solution is then added to the cell suspension in media or PBS at 1:1 volume ratio, for CCE formation, followed by pipette mixing and loading onto the flow cell. The light exposure time for macromonomer photo-initiated crosslinking was four seconds unless otherwise specified. After CCE formation, the remaining gel/cell mixture is washed with PBS or cell culture media, depending on the workflow.

To test the permeability of hydrogel CCEs (Supplementary Fig. 4), 10 minutes after wash, fluorescein dextran probes with 40, 150, 250, 500, or 2000 kDa molecular weights dissolved in PBS at 1 mg/mL concentration were loaded on flow cells and imaged on the instrument.

Fluorescein isothiocyanate–dextran probes with 40, 150, 250, and 2000 kDa molecular weights were purchased from Sigma. Dextran Fluorescein with 500 kDa molecular weight was purchased from Invitrogen.

To demonstrate the selective retention workflow (Fig. 2) an NK and Jurkat cell mix was prepared by mixing equal volumes of NK and Jurkat cell suspensions (each at 2 million cell/mL density) in a 1:1 volume ratio. After CCE formation using gel type A and washing, a solution of PE anti-human CD56 antibody (Biolegend, #362508) diluted 1:20 in PBS was loaded on the flow cell followed by 70 minutes of incubation at 37C and two washes with 200 uL PBS. To selectively compartmentalize the NK cells, gel type B was then loaded on the flow cell and larger secondary CCEs were formed around CD56-labelled cells followed by washing two times with 200 uL PBS. The primary CCEs (gel type A) were lysed by loading a solution of 5mM DTT in PBS.

### Optimization of CCE placement

We use two separate algorithms for calculating the optimal placement of CCEs, depending on whether the experiment uses a surface with capture spots (capture spot-aware CCE formation) or not (unconstrained CCE formation). For unconstrained CCE formation, we used simulated data to train a U-Net-like model(*71*) to predict, starting from the position of all the cells in the field of view, the likelihood that a cell can be compartmentalized, which generally depends on the local density of the cells. Once the model has identified the cells that can be compartmentalized, a CCE is placed, centered on the location of each cell. Overlaps between CCEs are resolved by moving CCEs apart using an iterative approximation algorithm, until a stopping criterion is met.

For capture spot-aware CCE formation we use a heuristic algorithm which calculates, for each cell, the number of “collisions” that would be incurred by placing a CCE over it. A collision is defined as an overlap with another CCE, or the overlap between a CCE and a capture spot (which would therefore not be available for occupation by another CCE). CCEs are then assigned in increasing number of collisions, under the constraint that no capture spot can be occupied by more than one CCE. Similar to the unconstrained case, overlaps between CCEs are resolved by an iterative approximation algorithm.

### Transcriptome assay

cDNA synthesis, followed by template switching to add a known handle at the 3^*′*^ end, is performed on the flow cell. Subsequently, the cDNA is eluted from the flow cell and PCR-amplified (15 cycles). The amplified product is bead-purified (0.6x SPRI), then fragmented and ligated with Illumina adapters using Watchmaker DNA library prep kit with fragmentation (#7K0022-096). The ligation product is bead-purified (0.8x SPRI), followed by another round of PCR during which sample indices are incorporated (8 cycles), and then bead-purified again (1x SPRI). The sequencing libraries were sequenced on the Illumina NextSeq 2000 platform.

### Standard Analysis and Quality Control of transcriptomic data

Sequencing data was processed using STARSolo(*72*) to generate a matrix of number of transcripts per gene per capture spot. The data is then aggregated at the CCE-level by summing the number of transcripts for each gene for all the capture spots associated with the same CCE. The CCE-level data is then filtered by only retaining CCEs that have a total number of transcripts above a threshold, a percent overlap with the oligo capture spot above a threshold, and a percentage of mitochondrial reads below a threshold. All the thresholds are user-defined and experiment-dependent. After these preprocessing and quality control steps, the data is processed in a standard Seurat(*73*) workflow comprising normalization, identification of highly variable features, scaling, PCA, selection of top PC dimensions, and UMAP embedding. All the data processing is implemented in an open-source R package, complete with documentation and example analysis vignettes. The R package also generates comprehensive quality control plots.

### Reproducibility and specificity of the transcriptome assay

To calculate gene expression signatures for individual cell lines (Fig. 3C), we performed differential gene expression analysis by comparing data derived from flow cell lanes loaded with individual cell lines. Specifically, we compared Raji cells against a combined dataset of NK-92 and TALL-104 cells, TALL-104 cells against NK-92 and Raji cells, and NK-92 cells against TALL-104 and Raji cells.

For each comparison, Seurat objects for the target and reference cell populations were merged, and a Seurat^42^ workflow was applied using SCTransform(*74*) to correct for unwanted sources of variation. Differentially expressed genes were identified using the Seurat *FindMarkers* function, selecting genes with an adjusted p-value ≤0.01 and a minimum log2 fold change of 5.

These differentially expressed genes were then used to calculate per-cell signature scores using the Seurat *AddModuleScore* function. The scores were scaled to the 0-1 range, and each cell within mixed lanes was assigned to the cell type with the largest cell type-specific signature score.

To assess specificity in the Barnyard experiment (Fig. 3D-E) the sequencing data was aligned on a combined reference comprising both the human and mouse genomes.

### Multi-modal profiling of PBMC following stimulation with LPS

Healthy human PBMCs were rested overnight after thawing (see “Tissue culture”), and then incubated with 50U/mL IL-2 for one hour. The cells were then stained with a 1:10 dilution of the Miltenyi TNF*α* catch antibody (Miltenyi, #130-091-267), 1:20 dilution of Alexa647-conjugated anti-CD14 antibody (Biolegend, #325611), and incubated with 10 µg/mL LPS for 75 minutes. Cells were then treated with 0.5 mM EDTA to prevent clumping, and resuspended in 10 µg/mL LPS in the presence of the PE-conjugated anti-TNF*α* Miltenyi detection antibody (Miltenyi, #130-091-267) at a 1:50 dilution and loaded on the instrument. Following CCE formation leftover hydrogel precursor was washed using media with LPS (5µg/mL) and Miltenyi detection antibody (1:50 dilution). The cells were incubated for a total of nine hours before lysis followed by transcriptome assay, for a total stimulation time of *∼*10 hours.

### Robustness of DINOv2 embeddings

To evaluate the robustness of DINOv2 embeddings under common imaging impairments, we conducted experiments using the embeddings extracted from single cell images and their perturbed versions from six adhered cell types: IMR-90, Hs675t Fibroblast, HeLa, MC38, mutuDC, and neuron soma. We adopted a classification-based strategy that reflects intra-class invariance. We trained a linear classifier (1024 nodes perceptron) to predict cell type labels directly from the DINOv2 embeddings of a clean (unperturbed) set of images dedicated to training (training set). We then compared the per-class accuracy of the classifier on clean and perturbed versions of the images from the set left out for testing (test set). This approach allows us to quantify the extent to which DINOv2 embeddings retain class-discriminative features under degradation, while implicitly accounting for morphological similarities within classes.

We considered several common impairments:

- Defocus blur: To simulate slight out-of-focus imaging, we leveraged multi-plane (z-stack) acquisitions of the same field of view. For each image, a random slice was selected from a range of focal offsets smaller than the depth of field of the objective
- Rotation: Random rotations of ±90^*°*^ or ±180^*°*^, simulating arbitrary cell orientations as the cells may be in any orientation when loaded on the flow cell
- Contrast and brightness shifts: Uniform shifts of ±20% in both brightness and contrast to model variability in imaging conditions.
- Gaussian blur: this was pplied with a kernel whose standard deviation was randomly sampled in the range [0, 2] pixels, simulating mild optical blur.
- Additive Gaussian noise: Pixel-wise independent Gaussian noise with a standard deviation randomly chosen from [0, 10] (in 8-bit intensity units 0-255), reflecting sensor noise and low-light conditions.
- Dropout noise: Random pixel dropout with a 5% probability, emulating dead pixels or small occlusions.

The classifier performance drop under each condition is plotted in Fig. 4B.

### Relating cell morphology to gene expression in Hs657.T fibroblasts

To cluster the data based on cell morphology features (Fig. 4E) we used the DINOv2 large model(*13*) to extract a 1024-dimensional feature vector for each cell. For 2% of the cases where a single CCE contained multiple cells, the feature vectors where averaged element-wise across all the cells in the CCE. The 1024-dimensional data was then embedded in a two-dimensional space using UMAP and clustered using k-means in the UMAP space. To identify differentially expressed genes between clusters (Fig. 4F) we used the *FindAllMarkers* function from the Seurat package and retained genes with an adjusted p-value ≤0.05.

### Profiling A549 cells following treatment with Olmutinib

A549 cells growing in culture were detached using Trypsin-EDTA (0.25%) (Fisher scientific) for 5 minutes at 37C to prepare a single-cell suspension, and labeled using Celltrace far red (ThermoFisher Scientific). The cell density was adjusted to 2e6 cells/mL before loading on a Fibronectin-coated flow cell. The flow cell was then placed in an incubator for 24 hours, after which the media was changed to include Annexin V-FITC (Biolegend), and the cells were imaged to acquire the baseline data. Three of the four lanes were then treated with media containing 10uM Olmutinib (MedChemExpress) and Annexin V-FITC, while the fourth lane did not contain any Olmutinib and was used as control. The media was refreshed every 8 hours for the following 32 hours and the cells were imaged after each media refresh. PE-Conjugated anti-EGFR (Biolegend) was added to the media prior to the last imaging scan. The cells were then lysed and processed for 3^*′*^ mRNAseq.

To define the phenotypic classes in Fig. 5A we first identified CCEs with segmented cells with an average area *<* 15000 pixels. This was necessary because the segmentation algorithm could not perfectly distinguish individual cell instances and would sometime merge multiple individual cells into a single detection. CCEs were then classified as follows: *daughter cell resistant* (CCEs with ≥1 dead cells and ≥1 live cells at the last timepoint); *cell cycle arrest* (CCEs containing exactly 1 live cell at all timepoints); *dead* (CCEs containing 0 live cells and ≥1 dead cells at the last timepoint). CCEs containing cells with pixel area *>* 15000, or containing ≥ 1 live cells at the last timepoint were classified as *proliferation*. 2,328 CCEs that could not be accurately classified were excluded from further analysis. Differential expression analysis was performed by using the *FindAllMarkers* function from the Seurat package. Significant genes were defined as having an adjusted p-value *<* 0.05 and an absolute log2 fold-change *>* 0.5. To identify pathway enrichments (Fig. 5E), we grouped CCEs by either their imaging-based classification, their gene expression cluster membership, or the cartesian product of the two. Only groups with ≥ 50 CCEs were retained to ensure statistical power. We downloaded MSigDB gene sets for human including Hallmark (H), Canonical Pathways C2:CP:BIOCARTA, C2:CP:PID, and Oncogenic (C6) collections. For each group in each grouping strategy, we created signed ranked gene lists using the formula: rank = -log10(p-value)×sign(log2FC). We performed fast preranked GSEA using *fgsea* and identified enriched pathways at FDR *<* 0.05 using Benjamini-Hochberg correction. To assess significant interaction effects between the different groupings (Fig. 5F) we fitted linear models with the formula: module_score ∼ RNA_cluster * imaging_phenotype, while correcting for lane number (as the treated CCEs are from three different lanes in the same flow cell) as a covariate in the model. We used as target modules the E2F, G2M, MYC, EMT, DNA REPAIR Hallmark gene sets from MSigDB, which were selected based on their biological relevance in this system. This approach identified pathways where the combination of RNA and imaging modalities revealed unique biology through significant interaction terms (adjusted p-value ≤0.05). To predict the four imaging-based phenotypic classes from the gene expression data we trained a multinomial elastic net model (*α*=0.5, 5-fold cross-validation) in R using the *glmnet* package. The model utilized the top 2000 variable genes as input features. We separated genes with non-zero coefficients by coefficient sign to identify positive and negative predictors for each phenotypic class. To characterize protein-protein interactions, we built STRING database networks (v12.0, medium confidence ≥ 400) for the top 50 positive and top 50 negative predictors for the *daughter cell resistance* class. For the survival analysis in TCGA patients, expression data and clinical annotations were accessed through cBioPortal(*75*) and the R2 Genomics Analysis and Visualization Platform (http://r2.amc.nl). We stratified patients by KCNMA1 and KCNN4 expression levels, comparing overall survival between the top quartile (highest 25% expression) and bottom quartile (lowest 25% expression). Kaplan-Meier survival curves were generated and statistical significance was assessed using log-rank tests.

### Preadipocyte differentiation

Human primary subcutaneous pre-adipocytes were obtained from ATCC (#PCS-210-010) and maintained in fibroblast basal media (ATCC, #PCS-201-030) (proliferation media) supplemented with Fibroblast Growth Kit low serum from (ATCC, #PCS-201-041). These cells were maintained in a 5% O2, 5% CO2, 37 ^*°*^C incubator. Adipogenesis was induced using Adipocytes Differentiation Toolkit for Adipose Derived MSCs and Preadipocytes (ATCC, # PCS-500-050). Upon cell detachment, 150.000 human primary preadipocytes were resuspended in 63 uL of proliferation media + 7uL of RGD and loaded on a fibronectin-coated flow cell. Upon CCE formation around individual cells, proliferation media was replaced with differentiation initiation media and the flow cell was incubated in a 5% O2, 5% CO2, 37 ^*°*^C incubator. Differentiation initiation media was replaced with differentiation maintenance media after 48 hours. The differentiation maintenance media was replaced manually every 24 hours up to day six. Brightfield images were collected along the differentiation induction at day 0, 1, 2, 5 and 6. On day 6 cells were stained with BODIPY 493/503 (MedChemExpress, #493-503), Image-IT™ LIVE Plasma Membrane and Nuclear Labeling Kit (ThermoFisher, #I34406) and imaged followed by lysis and 3^*′*^ mRNA sequencing.

### Microglia phagocytosis assay

BV2 microglia were dissociated and loaded on a fibronectin-coated flow cell. CCEs were formed around individual cells, in the presence of 50ug/mL of E. coli pHrodo Deep Red BioParticles (ThermoFisher, #P35360). Following CCE formation, the flow cell was incubated at 37C for 4h, to allow for cell adhesion and phagocytic activity, followed by imaging and lysis for 3^*′*^ mRNA sequencing.

### Analysis of preadipocyte differentiation and microglia phagocytosis data

We built elastic net models to predict lipid droplet accumulation by preadipocytes and the phagocytic activity of microglia cells. For both models we used as gene expression features all the genes that had non-zero expression in at least 15% of the CCEs. For the microglia dataset, we also used as image-derived features the 1024-dimensional DINOv2 embeddings of the individual cell crops. Imaging features were used either alone or in combination with gene expression features. For the preadipocyte data we used as response variable the ratio of the BODIPY lipid stain to the nuclear stain (Hoechst) across the entire CCE, as accurate segmentation of individual cell instances in brightfield was not possible. For the microglia dataset we used the log-transformed average pHrodo intensity across all the pixels belonging to each cell instance. We used 0.5 as the alpha value to mix L1 and L2 regularization penalties. The optimal lambda for the model was selected as the one resulting in the minimum average error over 10-fold cross-validation. Stable features were defined as those that appear in ≥ 50% of 100 bootstrap replicates. The model was implemented in R using the *glmnet* package. To identify the overlap with differentially expressed genes from transcriptomic clustering we used the FindAllMarkers function from Seurat with *min*.*pct*=0.25 *logfc*.*threshold*=0.5 and *only*.*pos=True*. The top-20 genes in each cluster (ordered by log2 fold-change) were selected.

### T cell priming assay

For the T cell priming assay (Supplementary Fig. 3) MutuDC cells were loaded on a fibronectin coated flow cell and incubated at 37C for 120 minutes for cells to adhere. Medium containing the SIINFEKL OVA-derived peptide (GenScript, #RP10611) at 1 µg/mL concentration was then flowed into the positive control lane and the cells were incubated for 60 minutes. OT1 cells mixed with hydrogel precursor were then flowed into the lane for CCE formation. The MutuDC express dsRed, which can be used to differentiate them from OT1, and were used as target cells for CCE formation. After CCE formation, the flow cell was loaded with medium containing an Alexa647-conjugated anti-CD69 antibody (Biolegend, #104517) at a 1:1000 dilution and the cells were imaged. Antibody-containing media at the same concentration was refreshed every six hours, followed by imaging.

## Acknowledgements

The authors wish to thank prof. Chris Mason and prof. Alexander Marson for helpful discussions.

## Authors contributions

TKK: Conceptualization, Methodology, Investigation, Writing; LYW: Conceptualization, Methodology, Investigation; PFG: Conceptualization, Software, Formal analysis, Writing; SM: Conceptualization, Methodology, Investigation; MM: Software, Formal analysis; FGY: Investigation; RD: Software, Methodology; JP: Methodology, Investigation; SS: Software, Formal analysis; AS: Investigation; YL: Investigation; NR: Investigation; NE: Investigation; SD: Formal analysis; AM: Formal analysis; RG: Investigation; SF: Investigation; IT: Investigation; CB: Methodology, Investigation; FC: Investigation; DR: Software; P-LH: Software, EZ-P: Investigation; RY: Investigation; OFV: Investigation; MP: Investigation; CX: Investigation; JW: Investigation; YZ: Investigation; PA: Investigation; MT: Investigation; EF: Investigation; BJ: Investigation; MS: Methodology; CV: Investigation; CC: Supervision; KGS: Methodology, Supervision; SJT: Methodology, Supervision; ORR: Supervision; FF: Supervision; JRE: Supervision; JRJ: Investigation; FHG: Supervision; MHS: Supervision, Writing; JP: Supervision, Writing; IK: Supervision; PL: Supervision; SL: Supervision, Writing; OO: Supervision, Funding acquisition; GPS: Supervision; MR: Supervision, Funding acquisition, Conceptualization, Writing.

