## Supplementary Figures for "Scalable longitudinal imaging and transcriptomics of cells in dynamic enclosures"

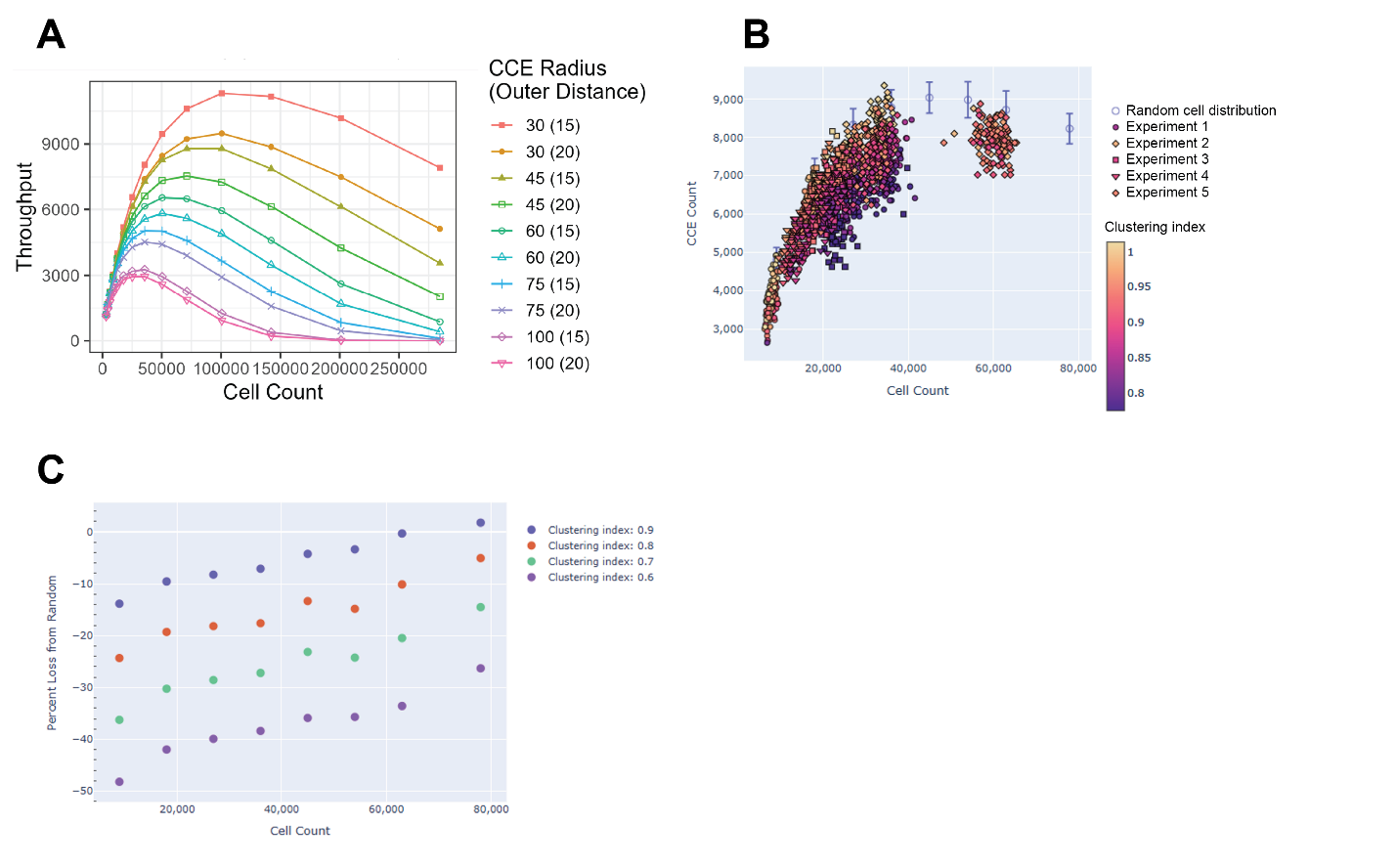

Supplementary Figure 1. Parameters affecting CCE formation throughput. A: Results from simulations relating the input cell density (x axis) and the CCE formation throughput, i.e. the number of CCEs formed in a flow cell lane (y axis). Each color and symbol combination corresponds to different settings of CCE radius (in µm) and the minimum distance to other CCEs (as displayed in the legend). Note that these simulations assume a uniform spatial distribution of cells. B: The theoretical throghput for a given combination of CCE formation settings is affected by the uniformity of cell loading (i.e. the amount of “clumpiness” in the sample). The open lilac circles represent the theoretical simulation under an assumption of uniform cell distribution. Each point in the plot indicates CCE formation throughput for a single FOV (extrapolated to the entire flow cell lane) for five different real experiments, indicated by the plot symbol. The color of the points correspond to a clustering index based on Quadrat Variance-To-Mean Ratio, indicating non-uniformity in the spatial dispersion of cells (with lower values indicating less uniformity or more “clumpiness”). C: simulations showing the percentage loss in CCE throughput for a simulation that assumes uniform spatial distribution of cells (y axis) vs sample density (x axis). Points of different colors correspond to simulations with different values of the clustering index.

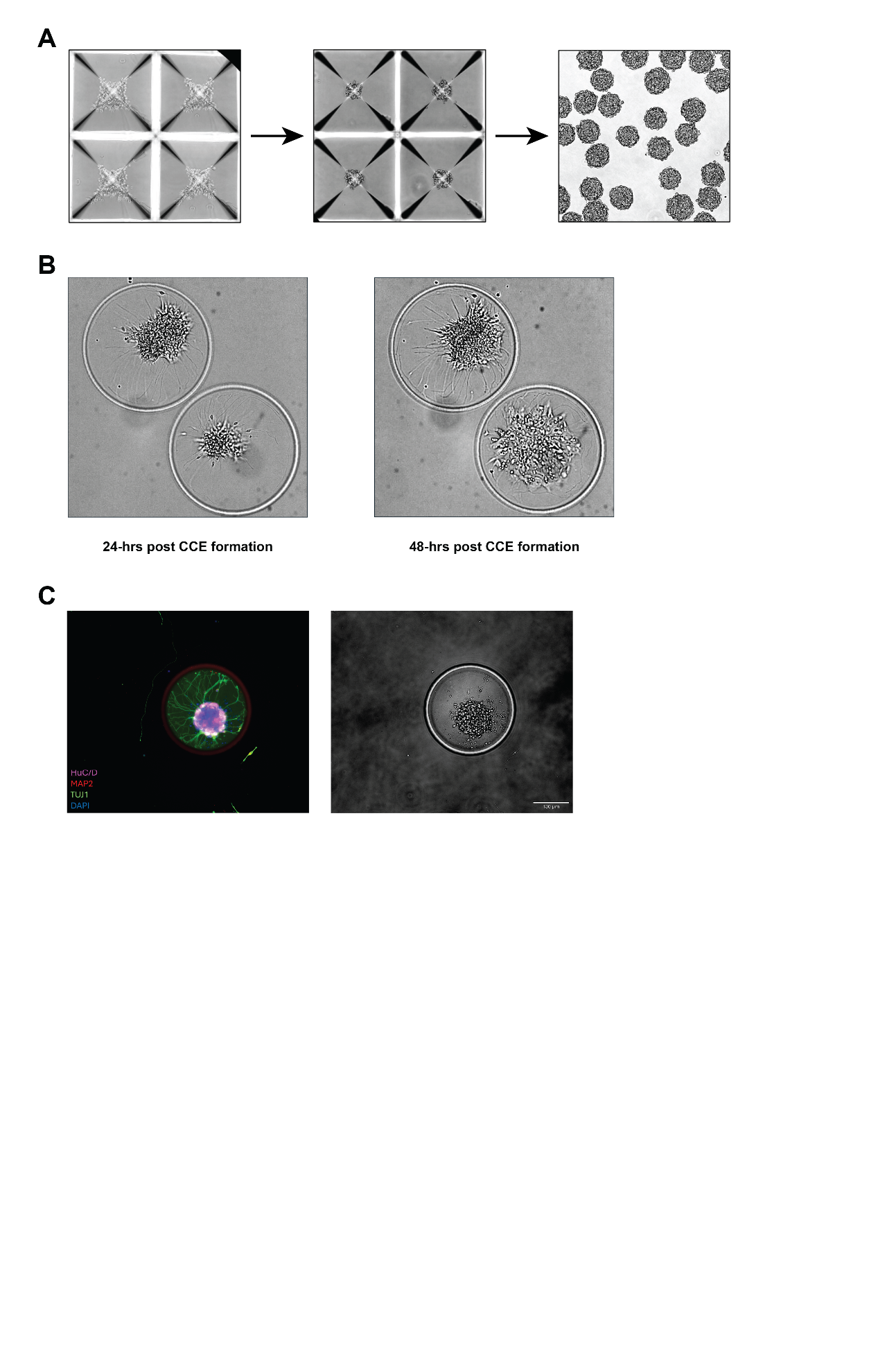

Supplementary Figure 2. Isolation of neurospheres in CCEs. A: neurospheres were generated from 19-day old iPS-derived V2a interneurons and 4-day old NGN2 induced neurons. Neurons were dissociated to single cells and then seeded into AggreWell plates to generate spheres of ~150 cells. After allowing cells to aggregate for 48 hours, they were passed through a 100 um strainer and loaded onto a flow cell. B: images of neurospheres 24 hours (left) and 48 hours (right) following CCE formation. C: immunofluorescence (left) and brightfield (right) imaging of a neurosphere, 48 hours after CCE formation. The immunofluorescence stains are, blue: DAPI, green: TUJ1, red: MAP2, magenta: HuC/D.

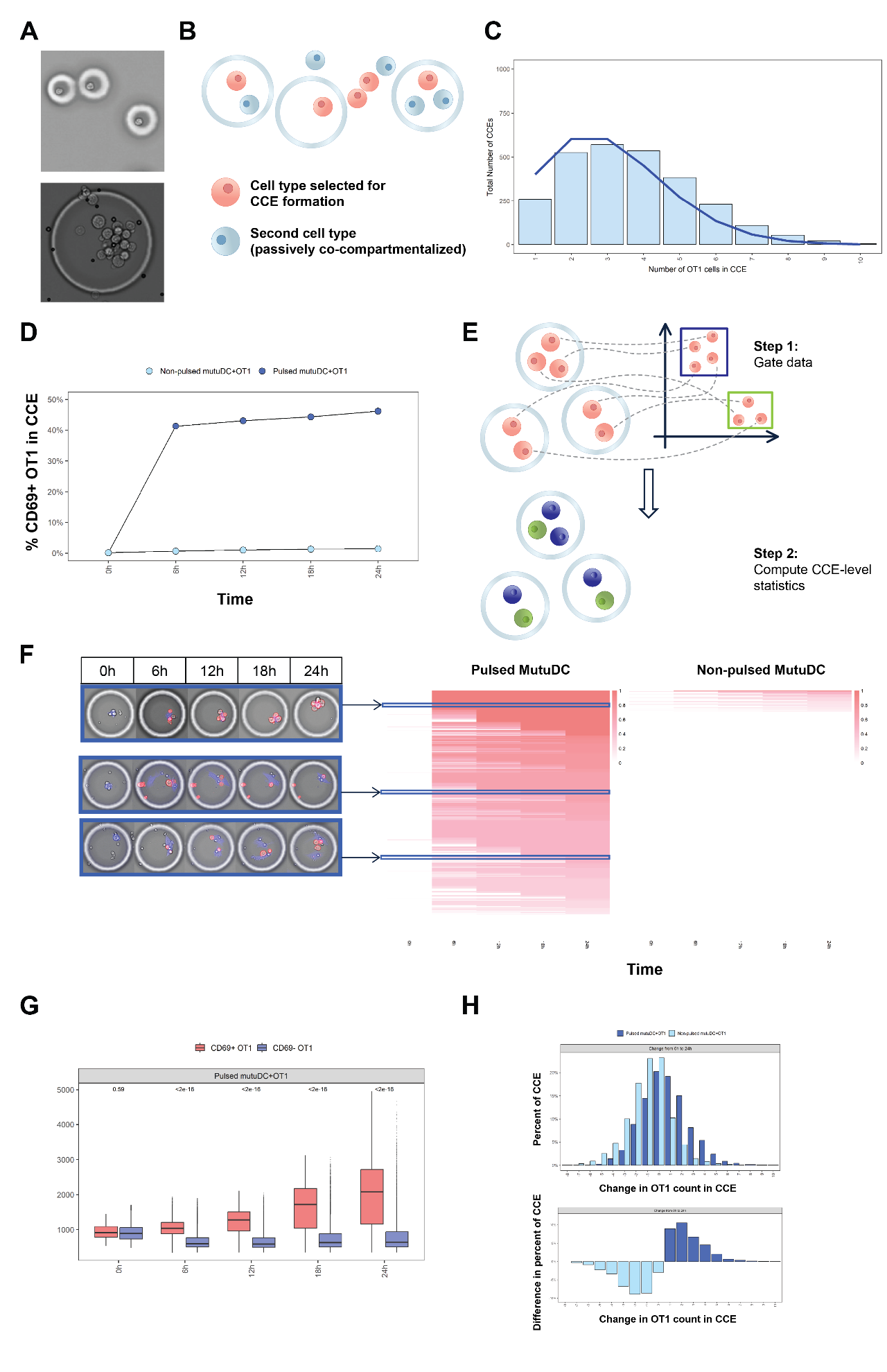
Supplementary Figure 3: T cell priming assay in CellCage™ Enclosures. The flexbile CCE formation logic enables the development of workflows aimed at studying cell-cell interactions, such as T cell priming assays. In these workflows one specific cell type is selected for compartmentalization, ensuring that each CCE contains an individual cell of interest. To ensure application of the correct compartmentalization logic during CCE formation, the two cell types are automatically identified and differentiated in real time by computer vision, e.g. by the presence or absence of different fluorescent stains, and a CCE is formed around the target single cell. The other cell type (termed “passenger”) is passively co-compartmentalized, resulting in a varying number of passenger cells per CCE that depends on their loading density. This produces a distribution of ratios of the two cell types, within each compartment. Across an entire flow cell lane this distribution closely mirrors the expectation under a Poisson model. To demonstrate the feasibility of this approach we developed a T cell priming assay using a murine dendritic cell line (MutuDC1940) and primary, antigen-specific, CD8+ T cells (OT-1), to assess the priming efficiency of individual dendritic cells. OT-1 cells express a transgenic T cell receptor that specifically recognizes the SIINFEKL peptide, derived from the Ovalbumin (OVA) protein, when presented by surface MHC class I molecules. MutuDCs pulsed with the peptide were seeded in the flow cell lane corresponding to the test condition. The negative control lane consisted of non-pulsed MutuDCs, which are not expected to prime the OVA-specific OT-1 cells. OT-1 cells mixed with hydrogel precursor were then flowed into the test and control lanes for CCE formation, and priming was monitored over 24 hours by staining for CD69.

A: small CCEs that contain individual cells (top) vs larger CCEs that contain multiple cells (bottom). B: CCE formation rules for T cell priming assay. MutuDC cells (red circles) are selected for CCE formation. OT-1 cells (blue circles) are passively co-compartmentalized, resulting in a varying number of OT-1 cells per CCE that depends on their density in the sample. C: distribution of the number of OT-1 cells across multiple CCEs (histogram), compared with the expectation under a Poisson model with λ =3 (blue line). D: Overall fraction of CD69+ OT-1 cells across all the CCEs (y axis) over time (x axis) in the lane where the MutuDCs were pulsed with the SIINFEKL peptide (dark blue) vs the negative control lane (non-pulsed, light blue), indicating higher T cell priming activity in the pulsed lane. E: analysis approach to generate CCE-level statistics. The cell data is pooled across the entire lane and gated to define populations of interest (e.g. MutuDC vs OT-1, CD69 positive vs negative etc.). The cell classification is then mapped back to individual CCEs to define CCE-level statistics (e.g. number of CD69+ OT-1 cells in the CCE). All the data displayed in this figure has been filtered to only include 5,080 CCEs with a single MutuDC and one or more OT-1 cells, following this classification process. F: heatmap showing heterogeneity in T cell priming activity (red color gradient) in individual CCEs (row) over time (columns). The T cell priming activity was defined, for each timepoint, as the maximum number of CD69+ OT-1 cells across all the timepoints up to the one of interest, divided by the number of OT-1 cells at that timepoint. At the end of the experiment, in the lane with antigen-pulsed MutuDCs, 69.2 % of the CCEs had a priming activity greater than 0.5, compared to only 1.5 % of CCEs in the control lane. The callouts display the imaging data associated with representative CCEs. G: boxplots displaying the distribution of cells sizes (y axis) over time (x axis) for CD69+ (red) vs CD69- (blue) OT-1 cells in the pulsed lane, indicating progression through the cell cycle of CD69+ OT-1 cells. H: the top histogram represents the distribution (y axis, as percentage) of the CCE-level differences in the number of OT-1 cells between the 24-hour timepoint and the baseline (x axis) in the pulsed (dark blue) vs non-pulsed (light blue) condition. 34.1% of the CCEs in the antigen-pulsed condition had an increase in the number of OT-1 cells, indicating proliferation following activation. The bar plot on the bottom represents the difference between the histograms for the two conditions in the top plot.

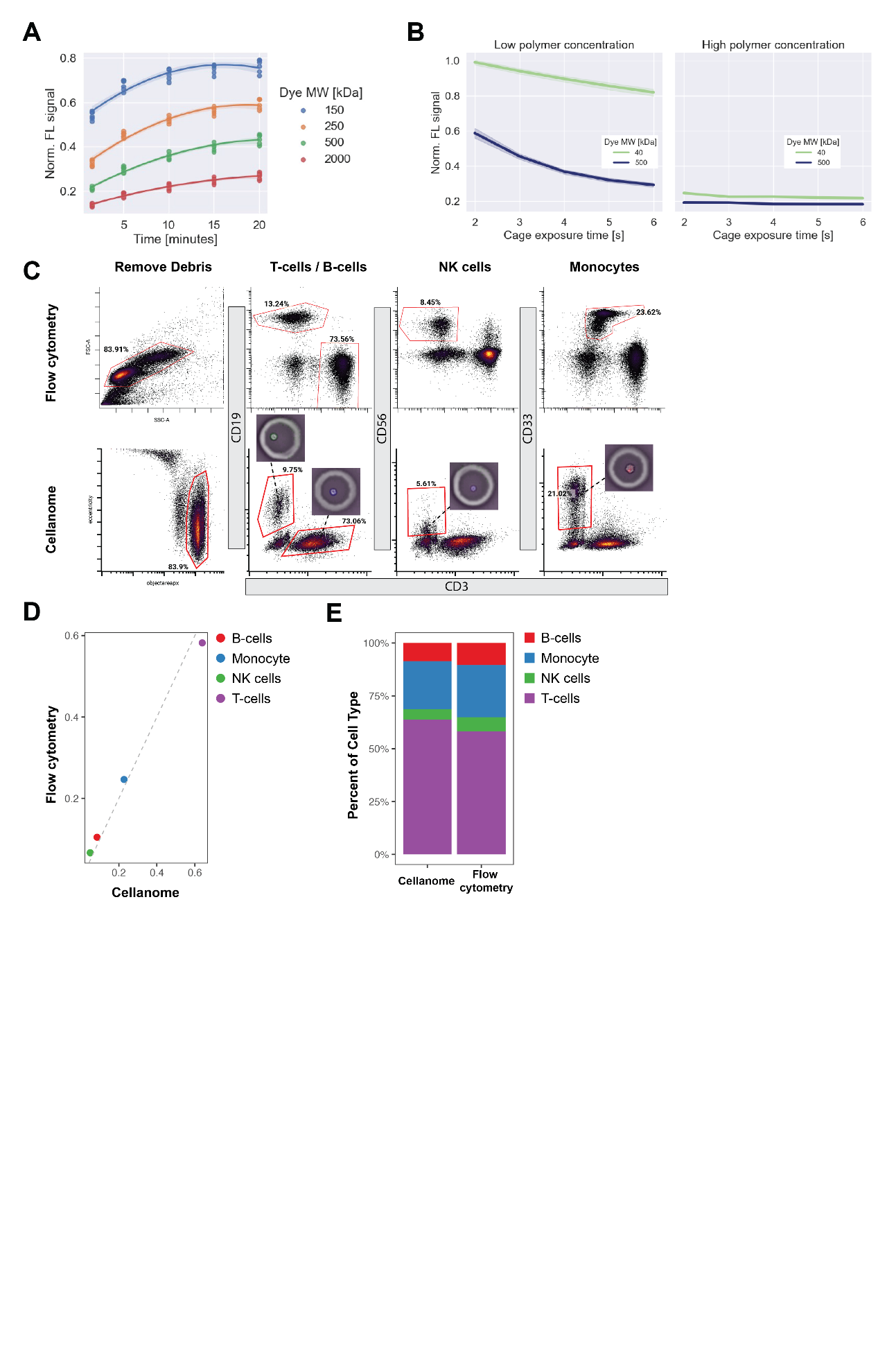

Supplementary Figure 4. Tunable porosity of the hydrogel material. A: permeability to dextran probes of various molecular weights (indicated by the line colors) measured as fluorescence signal in CCEs (y-axis) vs diffusion time (x-axis). B: permeability to dextran probes of various molecular weights (indicated by the line colors) measured as fluorescence signal in CCEs after 10 minutes of incubation at 37C (y-axis) as a function of CCE exposure time (x-axis). The left and right plot refer to low and high precursor polymer concentrations respectively. The permeability of the CCE walls enables antibody staining of compartmentalized cells. C: gating strategy for a four-marker surface receptor assay on healthy PBMCs performed on a conventional flow cytometer (top row) and on our platform (bottom row). The following antibodies were used

| Target | Clone | Vendor | Catalog No. | Fluorophore |
| --- | --- | --- | --- | --- |
| CD33 | P67.6 | Biolegend | 366625 | Alexa647 |
| CD3 | UCHT1 | Biolegend | 300406 | FITC |
| CD19 | HIB19 | Fisher Scientific | BDB753271 | RY586 |
| CD56 | HCD56 | Fisher Scientific | FAB2408V100 | Alexa405 |

For flow cytometry, the cells were stained for 30 minutes at room temperature in 100µl of cell staining buffer (Biolegend, #420201) with a 1:20 dilution of TruStainFcX block (Biolegend, #422302). All the antibodies were used at a 1:20 dilution. After washing, the cells were resuspended in cell staining buffer and the data was acquired on a Beckman Coulter CytoFlex instrument. For antibody staining within CCEs, the same mix was used and flowed in the lane after CCE formation around individual cells. Following incubation for 45 minutes, the lane was washed and imaged. D: scatterplot depicting the proportions of individual cell populations (identified by the colors) resulting from the gating in panel C as measured with conventional flow cytometry (y-axis) and with our platform (x-axis). The pictures correspond to representative single cells in CCEs that belong to each gate. E: stacked bar plots depicting the proportion of individual cell populations analyzed with our platform (left) or with conventional flow cytometry (right)

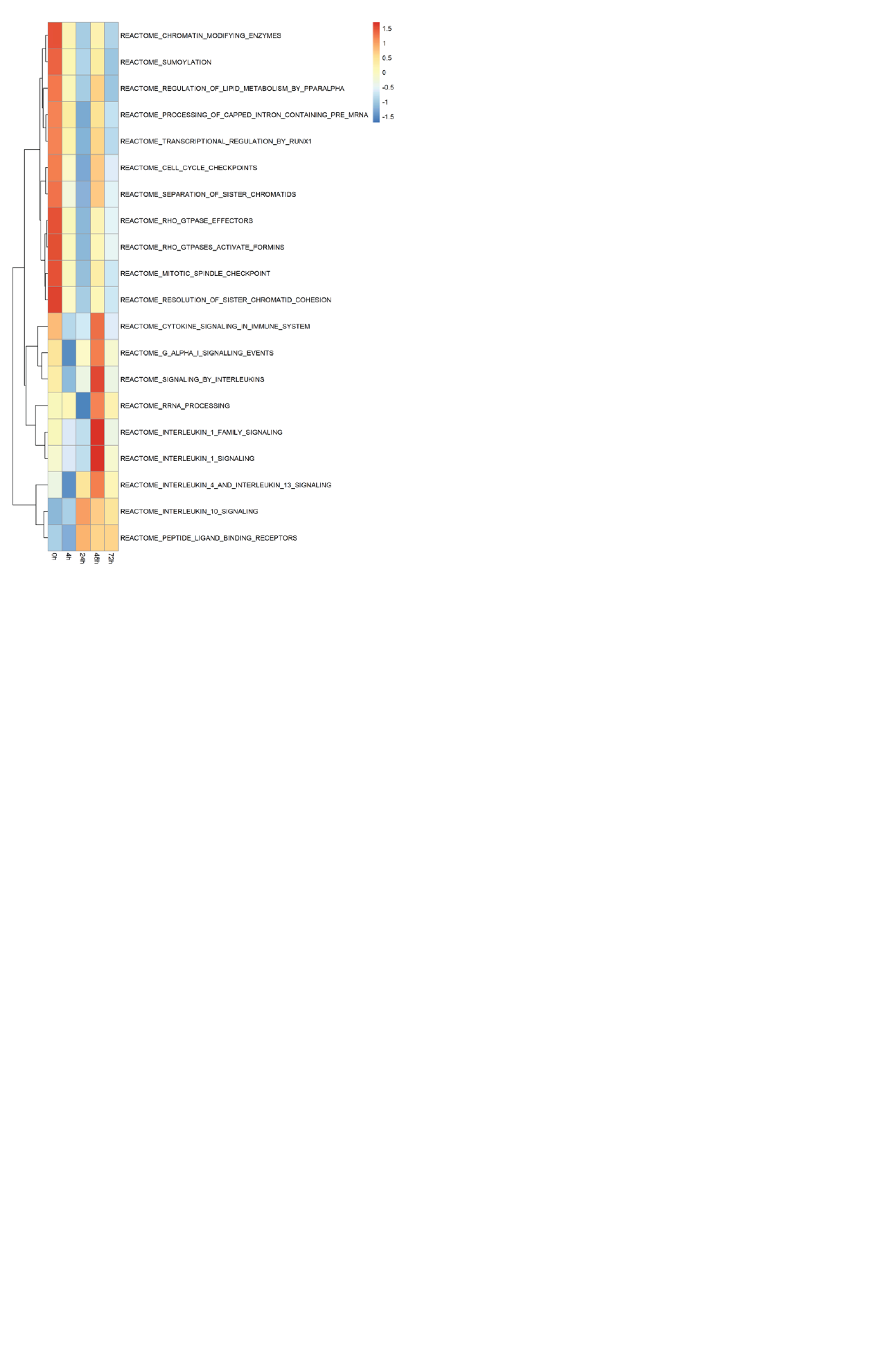

Supplementary Figure 5. Pathway activities in Hs675.T fibroblasts cultured on a fibronectin-coated flow cell. The heatmap represents pathway activation scores (rows) over time (columns) for all the Reactome pathways identified by Gene Set Enrichment Analysis of the individual differential expression results obtained by comparing successive timepoints (see Fig. 4D in the main text). Pathway activity scores are calculated as the average expression of all the genes in the pathway

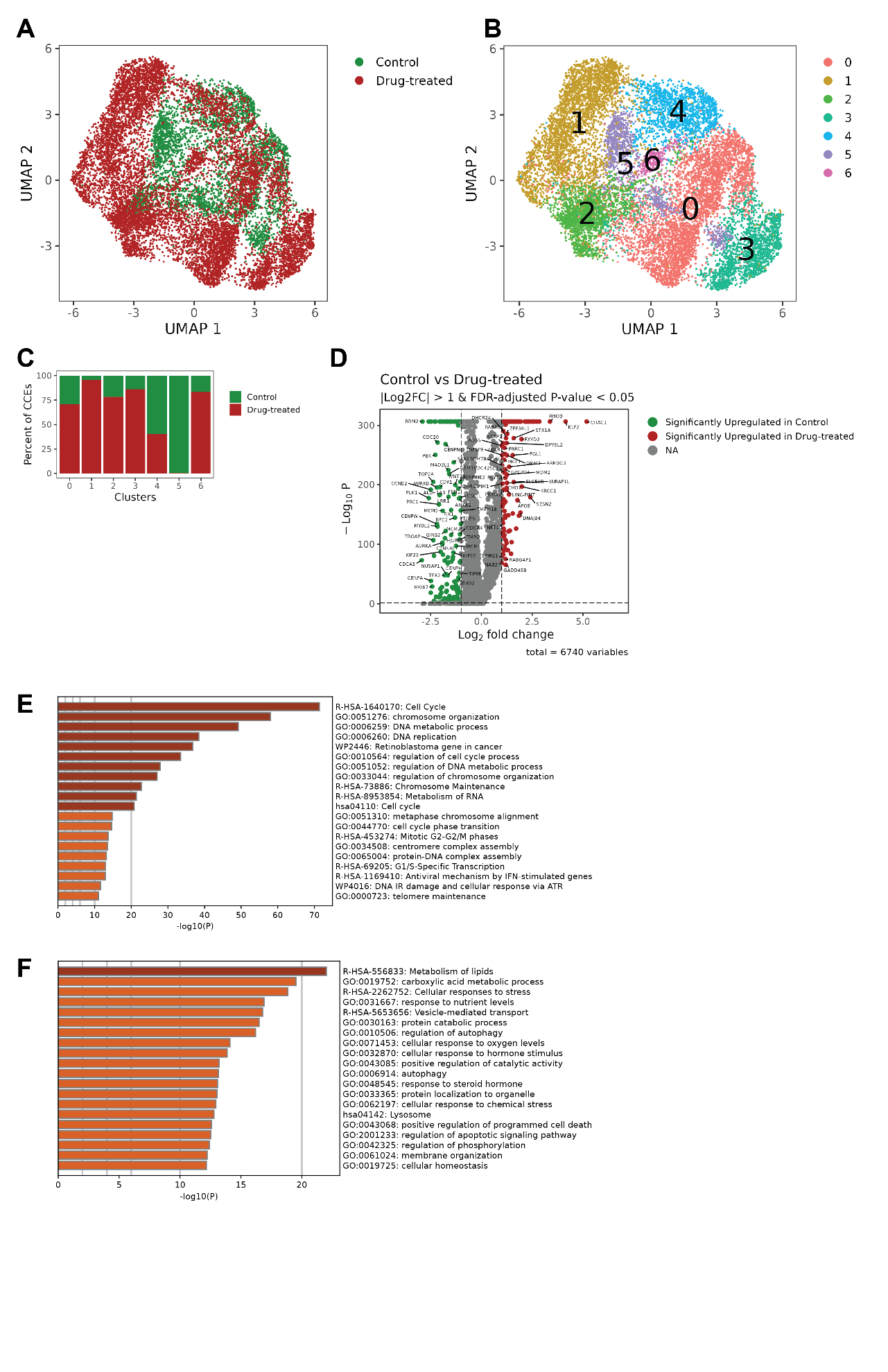

Supplementary Figure 6: Differential expression analysis between A549 cells treated with Olmutinib vs untreated. A: UMAP based on transcriptomic data. The experimental condition is indicated by the color. B: clustering of the gene expression data, the clusters are indicated by the color. C: proportion (y axis) of CCEs from each condition (indicated by the color) in each cluster (x axis). D: volcano plot representing the differential expression results between the two conditions (drug-treated vs control). E: Metascape enrichment results for the genes upregulated in the control condition. F: Metascape enrichment results for the genes upregulated in the drug-treated condition

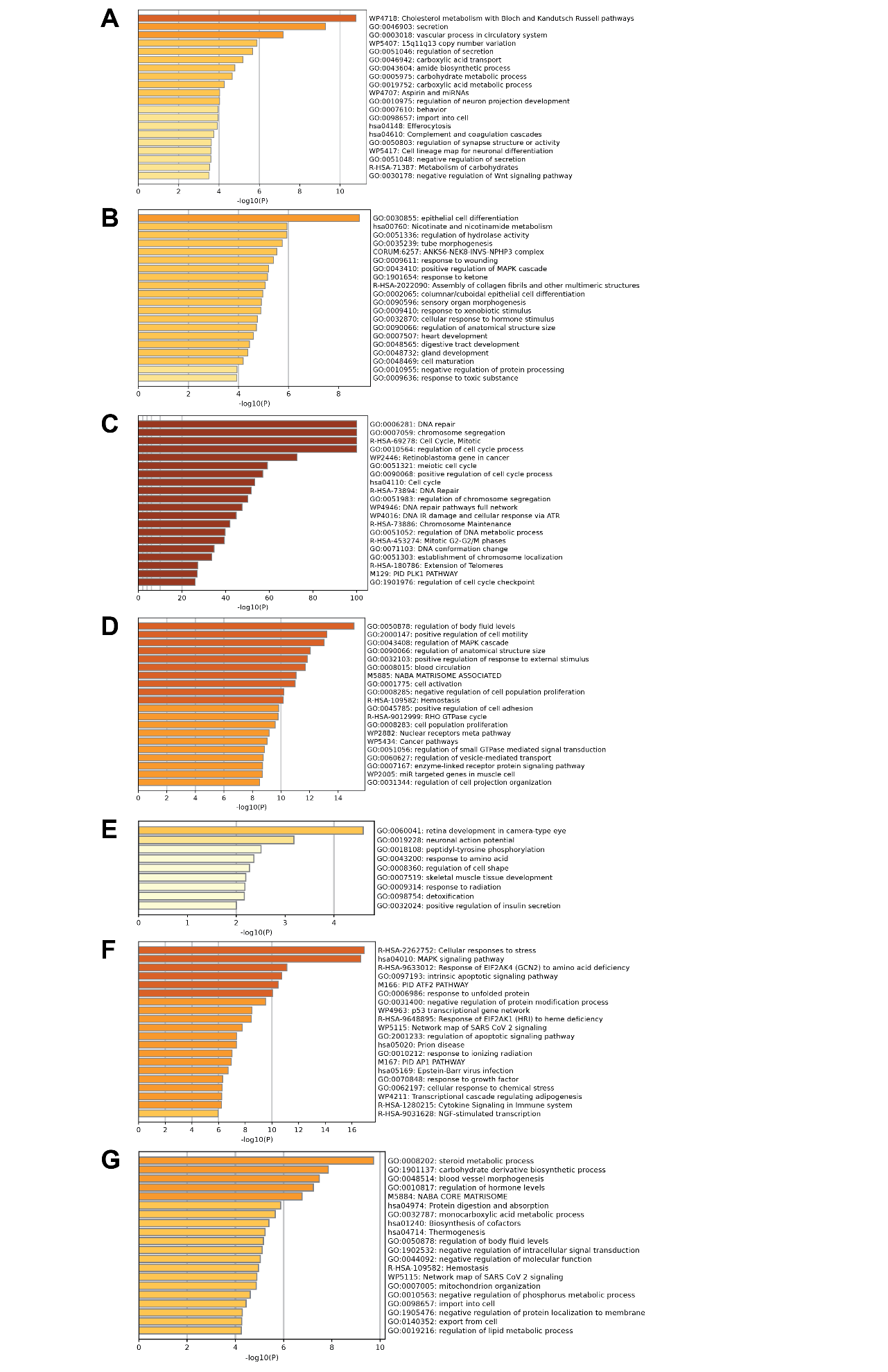

Supplementary Figure 7: Metascape enrichment results associated with the clustering of the transcriptomic data for A549 cells treated with Olmutinib. A-G: Metascape enrichment results for genes upregulated in clusters 0 through 6.
